# Non-invasive stimulation of the human striatum disrupts reinforcement learning of motor skills

**DOI:** 10.1101/2022.11.07.515477

**Authors:** Pierre Vassiliadis, Elena Beanato, Traian Popa, Fabienne Windel, Takuya Morishita, Esra Neufeld, Julie Duque, Gerard Derosiere, Maximilian J. Wessel, Friedhelm C. Hummel

## Abstract

Reinforcement feedback can improve motor learning, but the underlying brain mechanisms remain underexplored. Especially, the causal contribution of specific patterns of oscillatory activity within the human striatum is unknown. To address this question, we exploited an innovative, non-invasive deep brain stimulation technique called transcranial Temporal Interference Stimulation (tTIS) during reinforcement motor learning with concurrent neuroimaging, in a randomised, sham-controlled, double-blind study. Striatal tTIS applied at 80Hz, but not at 20Hz, abolished the benefits of reinforcement on motor learning. This effect was related to a selective modulation of neural activity within the striatum. Moreover, 80Hz, but not 20Hz tTIS increased the neuromodulatory influence of the striatum on frontal areas involved in reinforcement motor learning. These results show for the first time that tTIS can non-invasively and selectively modulate a striatal mechanism involved in reinforcement learning, opening new horizons for the study of causal relationships between deep brain structures and human behaviour.

## 1. Introduction

The ability to learn from past outcomes, often referred to as reinforcement learning, is fundamental for complex biological systems^1^. Reinforcement learning has been classically studied in the context of decision making, when agents have to decide between a discrete number of potential options^2^. Importantly, there is an increasing recognition that reinforcement learning processes are also at play in other contexts including during practice of a new motor skill^3–5^. For instance, the addition of reinforcement feedback during motor training can improve motor learning, presumably by boosting the retention of newly acquired motor memories^6,7^. Interestingly, reinforcement feedback also appears to be relevant for the rehabilitation of patients suffering from motor impairments^8–10^. Yet, despite these promising results, there is currently a limited understanding of the brain mechanisms that are critical to implement this behaviour.

A prominent hypothesis in the field is that the striatum, a structure that is particularly active both during reinforcement^11^ and motor learning^12^, may be causally involved in the beneficial effects of reinforcement on motor learning. As such, the striatum shares dense connexions with dopaminergic structures of the midbrain as well as with pre-frontal and motor cortical regions^13^, and is therefore well positioned to mediate reinforcement motor learning^14–16^. This idea is supported by neuroimaging studies showing reward-related activation of the striatum during motor learning^17,18^. More specifically, within the striatum, oscillatory activity in specific frequency bands is suggested to be involved in aspects of reinforcement processing. Previous rodent studies have shown that striatal high gamma oscillations (∼ 80 Hz) transiently increase following reward delivery^19–23^, but not when reward is withheld^19^. Hence, dynamic changes of high gamma activity in the striatum^19,24,25^ and in other parts of the basal ganglia^26,27^ may encode the outcome of previous movements (i.e., success or failure) and support learning. Consistent with a role of such oscillatory activity in reinforcement learning, high gamma activity in the striatum shows coherence with frontal cortex oscillations and is up-regulated by dopaminergic agonists^19^. Hence, this body of work suggests that reinforcement-related modulation of striatal oscillatory activity, especially in the gamma range, may be crucial for reinforcement learning of motor skills. Conversely, striatal beta oscillations (∼20 Hz) have been largely associated with sensorimotor functions^28^. For instance, beta oscillations in the striatum are exacerbated in Parkinson’s disease and associated to the severity of motor symptoms^29–31^. Consistently, excessive beta connectivity is reduced by anti-parkinsonian treatment in proportion to the related motor improvement^32^. Taken together, these elements suggest that striatal high gamma and beta activity may have different functional roles preferentially associated to reinforcement and sensorimotor functions, respectively.

The studies mentioned above provide associative evidence linking the presence of reinforcement with changes of neural activity within the striatum determined through neuroimaging^17,18^, but do not allow to draw conclusions regarding its causal role in reinforcement motor learning in humans. The only causal evidence available to date comes from animal work showing modulation of reinforcement-based decision-making with striatal stimulation^33,34^. A reason for the current absence of investigations of the causal role of the striatum in human behaviour is related to its deep localization in the brain. As such, current non-invasive brain stimulation techniques, such as transcranial magnetic stimulation (TMS) or classical transcranial electric stimulation (tES), do not allow to selectively target deep brain regions, because these techniques exhibit a steep depth-focality trade-off^35,36^. Studies of patients with striatal lesions^37,38^ or invasive deep brain stimulation of connected nuclei^39,40^ have provided insights into the role of the basal ganglia in reinforcement learning. However, their conclusions are partially limited by the fact that the studied patients also exhibit altered network properties resulting from the underlying pathology (e.g., neurodegeneration, lesions) or from the respective compensatory mechanisms. Here, we address these challenges by exploiting transcranial electric Temporal Interference Stimulation (tTIS), a new non-invasive brain stimulation approach allowing to target deep brain regions in a frequency-specific and focal manner in the physiological state^41,42^.

The concept of tTIS was initially proposed and validated on the hippocampus of rodents^41^ and was then further tested through computational modelling^43–47^ and in first applications on cortical areas in humans^48,49^. tTIS requires two pairs of electrodes to be placed on the head, each pair delivering a high frequency alternating current. One key element is that this frequency has to be high enough (i.e., in the kHz range) to avoid direct neuronal entrainment, based on the low-pass filtering properties of neuronal membranes^50^. The second key element is the application of a small difference of frequency between the two alternating currents. The superposition of the electric fields creates an envelope oscillating at this low-frequency difference, which can be steered towards individual deep brain structures (e.g., by optimizing electrodes’ placement), and is in a range able to influence neuronal activity ^41,51–53^. An interesting feature of tTIS is to stimulate at a particular frequency of interest in order to preferentially interact with specific neuronal processes^41,42^. Importantly, despite these exciting opportunities, current evidence for tTIS-related neuromodulation of deep brain structures, such as the striatum, is lacking in humans.

Here, we combine tTIS with electric field modelling for target localisation, behavioural data and functional magnetic resonance imaging (fMRI) to evaluate the causal role of specific patterns of striatal activity in reinforcement learning of motor skills. Based on the studies mentioned above, we hypothesised that striatal tTIS at high gamma frequency (tTIS_80Hz_) would disturb the fine-tuning of high gamma oscillatory activity in the striatum and thereby would perturb reinforcement motor learning in contrast to beta (tTIS_20Hz_) or sham (tTIS_Sham_) stimulation. More specifically, we reasoned that applying a constant high gamma rhythm in the striatum would disturb the temporally precise and reinforcement-specific modulation of high gamma activity. Moreover, given that the stimulation protocol was not individualised to endogenous high gamma activity and not synchronised to ongoing activity in other hubs of the reinforcement learning network (e.g., the frontal cortex), we anticipated disruptive rather than beneficial effects of tTIS_80Hz_.

In line with our prediction, we report that tTIS_80Hz_ disrupted motor learning compared to the controls, but only in the presence of reinforcement. To evaluate the potential neural correlates of these behavioral effects, we measured BOLD activity in the striatum and effective connectivity between the striatum and frontal cortical areas involved in reinforcement motor learning. We found that the disruptive effect of tTIS_80Hz_ on reinforcement learning was associated to a specific modulation of BOLD activity in the putamen and caudate, but not in the cortex, supporting the ability of tTIS to selectively modulate striatal activity without affecting overlying cortical areas. Moreover, tTIS_80Hz_ also increased the neuromodulatory influence of the striatum on frontal cortical areas involved in reinforcement motor learning. Overall, the present study shows for the first time that tTIS can non-invasively and selectively modulate a striatal mechanism involved in reinforcement learning opening new horizons for the study of causal relationships between deep brain structures and human behaviour.

## 2. Results

24 healthy participants (15 women, 25.3 ± 0.1 years old; mean ± SE) performed a force tracking task in the MRI with concurrent tTIS of the striatum. The task required participants to modulate the force applied on a hand-grip force sensor in order to track a moving target with a cursor with the right, dominant hand^54,55^ (**Figure 1A**). At each block, participants had to learn a new pattern of motion of the target (**Figure S1**; see Methods). In Reinf_ON_ blocks, participants were provided with online reinforcement feedback during training, giving them real-time information about success or failure throughout the trial, indicated as a green or red target, respectively (please see **Video S1** for the task). The reinforcement feedback was delivered according to a closed-loop schedule^8^, in which the success criterion to consider a force sample as successful was updated based on the median performance over the 4 previous trials (see Methods for more details). In Reinf_OFF_ blocks, participants practiced with a visually matched random feedback (cyan/magenta). Importantly, in both types of blocks, training was performed with partial visual feedback of the cursor, a condition that has been shown to maximise reinforcement effects in various motor learning paradigms^4,56–58^ and which yielded significant effects of reinforcement on motor learning as also demonstrated in an additional behavioural study testing another group of healthy participants on the same task (n = 24, Figure S2). Before and after training, participants performed Pre- and Post-training assessments with full visual feedback, no reinforcement and no tTIS, allowing us to evaluate motor learning. To assess the effect of tTIS on reinforcement-related benefits in motor learning and the associated neural changes, participants performed 6 blocks of 36 trials in the MRI, with concurrent tTIS during training, delivered with a Δf of 20 Hz (tTIS_20Hz_), 80 Hz (tTIS_80Hz_) or as a sham (tTIS_Sham_; 3 tTIS_TYPE_ x 2 Reinf_TYPE_ conditions; **Figure 1B, 1C**). Notably, the order of the conditions was balanced among the 24 participants, ensuring that any potential carry-over effect would have the same impact on each experimental condition. To determine the best electrode montage to stimulate the human striatum (putamen, caudate and nucleus accumbens [NAc] bilaterally), computational modelling with a realistic head model was conducted with Sim4Life^59^ (see Methods). The selected montage (F3-F4; TP7-TP8) generated a theoretical temporal interference electric field that was ∼30-40% stronger in the striatum than in the overlying cortex, reaching magnitudes of 0.5 to 0.6 V/m (**Figure 1D, 1E**).

**Figure 1.**
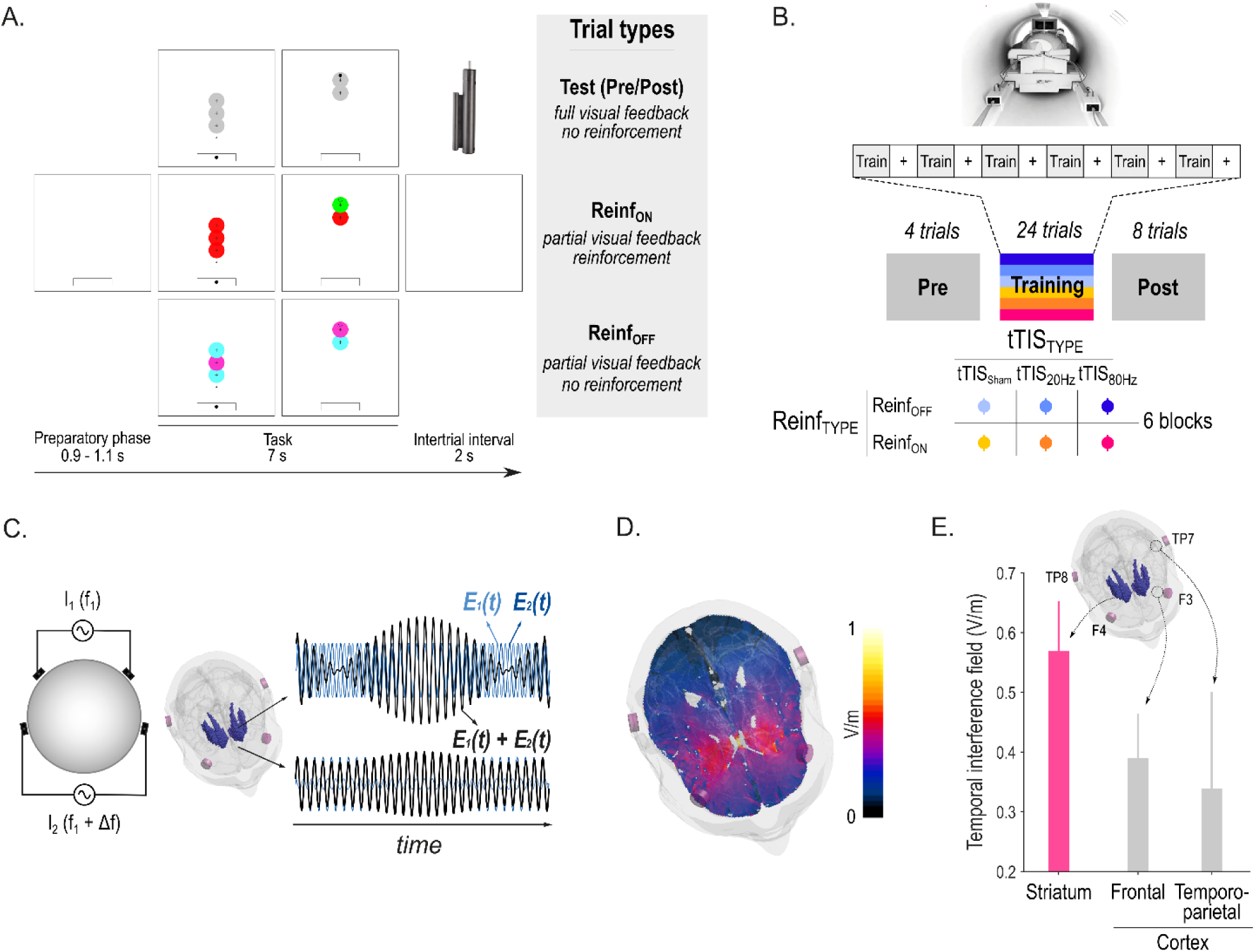
Striatal tTIS during reinforcement learning of motor skills in the MRI. **A) Motor learning task.** Participants were required to squeeze a hand grip force sensor (depicted in the upper right corner of the figure) in order to track a moving target (larger circle with a cross in the center) with a cursor (black smaller circle)^54,55^. Pre- and Post-training assessments were performed with full visual feedback of the cursor and no reinforcement. In Reinf_ON_ and Reinf_OFF_ trials, participants practiced the task with or without reinforcement feedback, respectively. As such, in Reinf_ON_ trials, the color of the target varied in real-time as a function of the subjects’ tracking performance. **B) Experimental procedure.** Participants performed the task in the MRI with concomitant TI stimulation. Blocks of training were composed of 36 trials (4 Pre-, 24 Training and 8 Post-training trials) interspersed with short resting periods (represented as + on the figure). The 6 training types resulted from the combination of 3 tTIS_TYPES_ and 2 Reinf_TYPES_. **C) Concept of tTIS.** On the left, two pairs of electrodes are shown on a head model and currents are applied with a frequency f1 and f1+Δf. On the right, the interference of the two electric fields within the brain is represented for two different locations with respectively high and low envelope modulation. E_1_(t) and E_2_(t) represent the modulation of the fields’ magnitude over time. tTIS was delivered either with a Δf of 20 or 80 Hz or as a sham (ramp-up and immediate ramp-down of high frequency currents with flat envelope). **D) Electric field modelling with the striatal montage.** Temporal interference exposure (electric field modulation magnitude). **E) Temporal interference exposure averaged in the striatum and in the overlying cortex.** Magnitude of the field in the cortex was extracted from the Brainnetome atlas (BNA^60^) regions underneath the stimulation electrodes (F3-F4 and TP7-TP8). Error bars represent the standard deviation over the voxels in the considered region.

### tTIS_80Hz_ disrupts reinforcement learning of motor skills

Task performance was evaluated by means of the Error, which was defined as the absolute difference between the applied and target force averaged across samples for each trial, as done previously^4,54,56^ (**Figure 2A**). Across conditions, the Post-training Error was reduced compared to the Pre-training Error (single sample t-test on the normalised Post-training data: t_(24)_=-2.69; p=0.013; Cohen’s d=-0.55), indicating significant motor learning during the task (**Figure 2B**). Such improvement was greater when participants had trained with reinforcement (Reinf_TYPE_ effect in the Linear Mixed Model (LMM): F_(1, 1062.2)_=5.17; p=0.023; d=-0.14 for the post-hoc contrast Reinf_ON_ – Reinf_OFF_), confirming the beneficial effect of reinforcement on motor learning^7,58^. Crucially though, this effect depended on the type of stimulation applied during training (Reinf_TYPE_ x tTIS_TYPE_ interaction: F_(2, 1063.5)_=2.11; p=0.034; **Figure 2C**). While reinforcement significantly improved learning when training was performed with tTIS_Sham_ (p=0.036; d=-0.22) and tTIS_20Hz_ (p=0.0089; d=-0.27), this was not the case with tTIS_80Hz_ (p=0.43; d=0.083). Consistently, direct between-condition comparisons showed that in the Reinf_ON_ condition, learning was reduced with tTIS_80Hz_ compared to tTIS_20Hz_ (p=0.039; d=0.26) and tTIS_Sham_ (p<0.001; d=0.45) but was not different between tTIS_20Hz_ and tTIS_Sham_ (p=0.15; d=0.20). This disruption of motor learning with tTIS_80Hz_ was not observed in the absence of reinforcement (tTIS_80Hz_ vs. tTIS_20Hz_: p=0.59; d=-0.10, tTIS_80Hz_ vs. tTIS_Sham_: p=0.34; d=0.15). These results strongly point to the fact that high gamma striatal tTIS specifically disrupts the benefits of reinforcement on motor learning and not motor learning in general.

**Figure 2.**
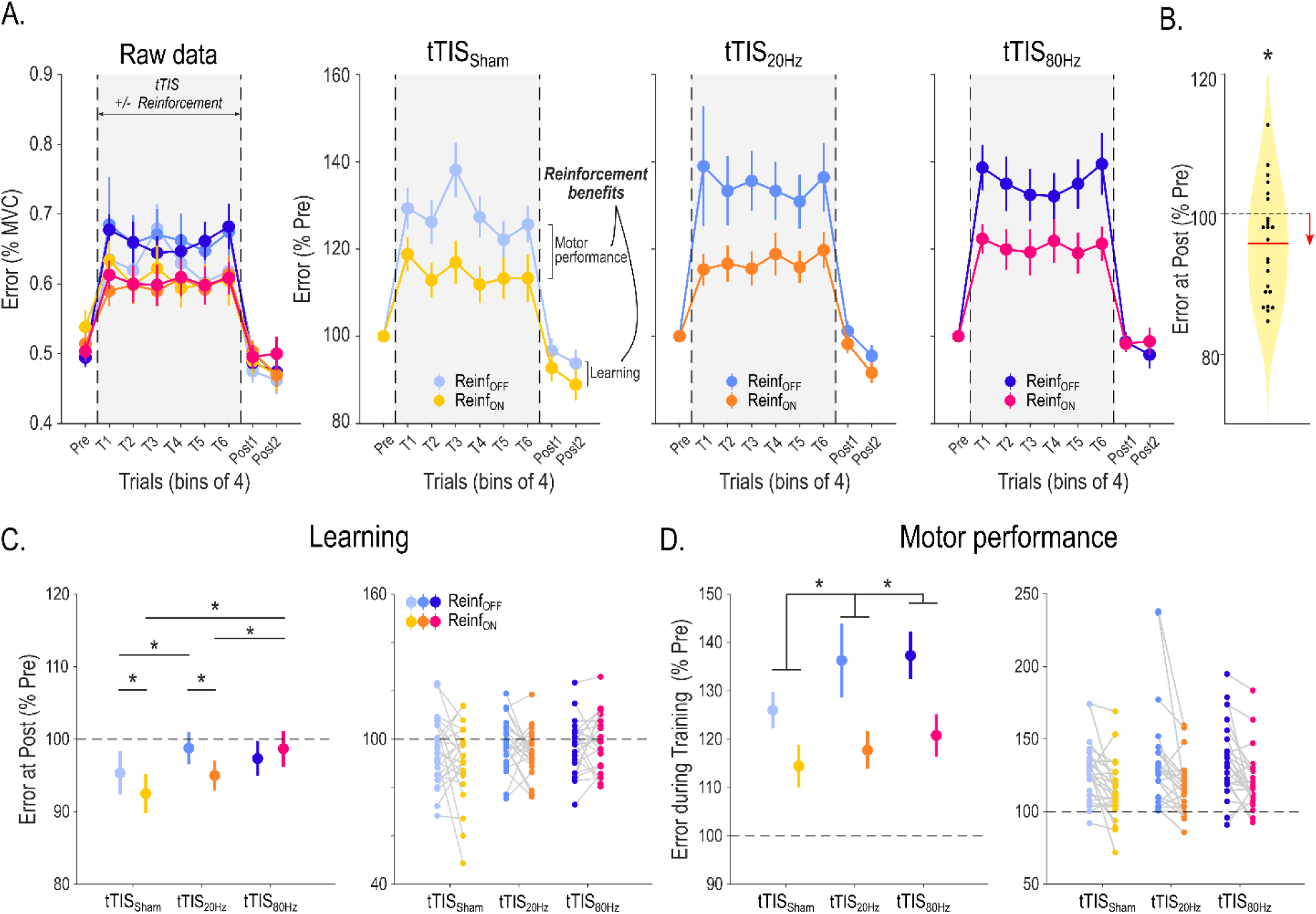
Behavioural results. **A) Motor performance across training.** Raw Error data (expressed in % of Maximum Voluntary Contraction [MVC]) are presented on the left panel for the different experimental conditions in bins of 4 trials. The increase in Error during Training is related to the visual uncertainty (i.e., intermittent disappearance of the cursor) that was applied to enhance reinforcement effects. On the right, the three plots represent the Pre-training normalised Error in the tTIS_Sham,_ tTIS_20Hz_ and tTIS_80Hz_ blocks. Reinforcement-related benefits represent the improvement in the Error measured in the Reinf_ON_ and Reinf_OFF_ blocks, during Training (reflecting benefits in motor performance) or at Post-training (reflecting benefits in learning). **B) Averaged learning across conditions.** Violin plot showing the Error distribution at Post-training (expressed in % of Pre-training) averaged across conditions, as well as individual subject data. A single-sample t-test showed that the Post-training Error was reduced compared to the Pre-training level, indicating significant learning in the task. **C) Motor learning.** Averaged Error at Post-training (normalised to Pre-training) and the corresponding individual data points in the different experimental conditions are shown on the left and right panels, respectively, for the subjects included in the analysis (i.e., after outlier detection, remaining n=23). Reduction of Error at Post-training reflects true improvement at tracking the target in Test conditions (in the absence of reinforcement, visual uncertainty or tTIS). The LMM ran on these data revealed a specific effect of tTIS_80Hz_ on reinforcement-related benefits in learning. **D) Motor performance.** Averaged Error during Training (normalised to Pre-training) and the corresponding individual data points in the different experimental conditions are shown on the left and right panels, respectively, for the subjects included in the analysis (i.e., after outlier detection, n=23). Individual data points are shown on the right panel. Error change during Training reflect the joint contribution of the experimental manipulations (visual uncertainty, potential reinforcement and tTIS) on motor performance. The LMM ran on these data showed a frequency-dependent effect of tTIS on motor performance, irrespective of reinforcement. *: p<0.05. Data are represented as mean ± SE.

Although training with tTIS_20Hz_ did not alter the benefits of reinforcement on motor learning, we found that learning without reinforcement was significantly impaired in this condition (tTIS_20Hz_ vs. tTIS_Sham_: p=0.046; d=0.25, **Figure 2C**). This suggests that tTIS_20Hz_ may disrupt a qualitatively different mechanism involved in motor learning from sensory feedback^61^, in line with the role of striatal beta oscillations in sensorimotor function^28^.

Next, we evaluated the effect of tTIS on motor performance during training itself. As shown in Figure 2A, the Error was generally higher during Training than in Test trials due to the presence of visual uncertainty during this phase. The extent of this disruption was reduced in the presence of reinforcement (Reinf_TYPE_: F_(1, 3262.4)_=339.89; p<0.001; d=-0.64 for the contrast Reinf_ON_ – Reinf_OFF_), demonstrating the ability of subjects to exploit real-time reinforcement information to improve tracking (**Figure 2D**). Notably, this effect was not modulated by tTIS_TYPE_ (Reinf_TYPE_ x tTIS_TYPE_: F_(2, 3265.8)_=0.91; p=0.40), indicating that tTIS did not directly influence reinforcement gains during tracking. Interestingly though, striatal stimulation did impact on general tracking performance independently of reinforcement as indicated by a significant tTIS_TYPE_ effect (tTIS_TYPE_: F_(2, 3262.4)_=42.85; p<0.001). This effect was due to an increase in the Error when tTIS_20Hz_ was applied (p<0.001; d=0.28 when compared to tTIS_Sham_), which was even stronger during tTIS_80Hz_ (p<0.001; d=0.38 and p=0.031; d=0.11 when compared to tTIS_Sham_ and tTIS_20Hz_, respectively). An additional analysis showed that the detrimental effect of tTIS on motor performance was actually due to an impaired ability to improve performance during Training (LMM with continuous fixed effect Trial: tTIS_TYPE_ x Trial interaction: F_(2, 3399)_=4.46; p=0.012, post-hoc tests: tTIS_Sham_ vs. tTIS_20Hz_: p=0.013; tTIS_Sham_ vs. tTIS_80Hz_: p=0.068; tTIS_20Hz_ vs. tTIS_80Hz_: p=0.81; **Figure S3**). However, again, this effect did not depend on the presence of reinforcement (Reinf_TYPE_ x tTIS_TYPE_ x Trial: F_(2, 3399)_=0.51; p=0.60). Notably, we also found that the detrimental effect of striatal tTIS did not depend on the availability of visual information on the cursor, but rather that tTIS had a general effect on motor performance irrespective of visual and reinforcement feedback (see Supplementary materials). This analysis also confirmed that reinforcement gains in motor performance were stronger when visual information was not available (**Figure S4**), in line with the behavioural data mentioned above (Figure S2) and previous studies^57,62^. Overall, these results suggest that striatal tTIS altered motor performance in a frequency-dependent manner but did not influence the ability to rapidly adjust motor commands based on reinforcement feedback during training. Hence, tTIS_80Hz_ may not disrupt real-time processing of reinforcement feedback, but may rather impair the beneficial effect of reinforcements on the retention of motor memories^6,7^.

Notably, these effects could not be explained by potential differences in initial performance between conditions (Reinf_TYPE_ x tTIS_TYPE_: F_(2, 519.99)_=1.08; p=0.34), nor by changes in the flashing properties of the reinforcement feedback (i.e., the frequency of color change during tracking; Reinf_TYPE_ x tTIS_TYPE_: F_(2, 3283)_=0.19; p=0.82), or by differences in success rate in the Reinf_ON_ blocks (i.e., the proportion of success feedback during tracking; tTIS_TYPE_: F_(2, 1702)_=0.17; p=0.84). The Reinf_TYPE_ x tTIS_TYPE_ effect on learning was also not influenced by the order of the reinforcement conditions (analysis on sub-groups based on whether participants experienced Reinf_ON_ or Reinf_OFF_ first; no Reinf_TYPE_ x tTIS_TYPE_ x Group_TYPE_ interaction: F(2,1105.06)=1.75; p=0.17; see Supplementary materials for more details on these analyses).

Finally, we confirmed that these results were not a consequence of an inefficient blinding. As such, when debriefing after the experiment, only 6/24 participants were able to successfully identify the order of the stimulation applied (e.g., real – real – placebo; chance level: 4/24; Fisher exact test on proportions: p=0.74). Consistently, the magnitude (**Figure S5A**) and type (**Figure S5B**) of tTIS-evoked sensations evaluated before the experiment were qualitatively similar across conditions and tTIS was generally well tolerated in all participants (no adverse events reported). This suggests that blinding was successful and is unlikely to explain our findings. More generally, this is a first indication that tTIS evokes very limited sensations (e.g., only 2/24 and 1/24 subjects rated sensations evoked at 2 mA as “strong” for tTIS_20Hz_ and tTIS_80Hz_, respectively; **Figure S5A**) that are compatible with efficient blinding.

### The effect of tTIS_80Hz_ on reinforcement motor learning is related to modulation of neural activity in the striatum

As mentioned above, task-based fMRI was acquired during Training with concomitant tTIS. This allowed us to evaluate the neural effects of tTIS and their potential relationship to the behavioural effects reported above. As a first qualitative evaluation of the data, we performed a whole-brain analysis in the tTIS_Sham_ condition to assess the network activated during reinforcement motor learning (Reinf_ON_ condition). Consistent with previous neuroimaging studies employing similar tasks^63,64^, we found prominent BOLD activations in a motor network including the putamen, thalamus, cerebellum and sensorimotor cortex, particularly on the left hemisphere, contralateral to the trained hand (**Figure S6, Table S2**). Notably though, contrasting Reinf_ON_ and Reinf_OFF_ conditions did not reveal any significant cluster at the whole-brain level. Hence, this first analysis did not reveal any region specifically activated in the presence of reinforcement, but rather confirms the involvement of a motor network engaged in this type of task irrespective of the reinforcement feedback.

As a second step, we evaluated the effect of tTIS on striatal activity, as a function of the type of reinforcement feedback and focusing on the very same regions of interest (ROI) that were used to optimise tTIS exposure in the modelling. Based on this, we extracted averaged BOLD activity within the bilateral putamen, caudate and NAc based on the Brainnetome atlas (BNA^60^), in the different experimental conditions and considered these six striatal ROIs (ROI_STR_) as fixed effects in the LMM. This model revealed a strong enhancement of striatal activity with Reinf_ON_ with respect to Reinf_OFF_ (F_(1, 800.01)_=13.23; p<0.001; d=0.25 for the contrast Reinf_ON_ – Reinf_OFF_) consistent with previous literature^11^, but no tTIS_TYPE_ effect (F_(2, 800.01)_=0.46; p=0.63) and no interaction (all p> 0.65; **Figure 3A**). Despite the absence of effects of tTIS on averaged striatal activity, we then asked whether the behavioural effects of tTIS_80Hz_ on reinforcement motor learning (i.e., tTIS_80Hz_ vs. tTIS_20Hz_ and tTIS_Sham_ with Reinf_ON_) could be linked to modulation of activity in core brain regions. To do so, we ran a whole-brain analysis focusing on the main behavioural effects mentioned above. Results revealed that the effect of tTIS_80Hz_ (with respect to tTIS_20Hz_) on motor learning in the Reinf_ON_ condition was specifically related to modulation of activity in two clusters encompassing the left putamen and bilateral caudate (**Figure 3B, Table S3**). Notably, the presence of the high frequency carrier (kHz) in both stimulation conditions rules out the possibility that the correlation was due to putative neuromodulatory effects of high frequency stimulation. No significant clusters were found neither for the tTIS_80Hz_ – tTIS_Sham_ contrast, nor for the control tTIS_20Hz_ -tTIS_Sham_ contrast, indicating that the reported correlation is not due to a general link between striatal activity and reinforcement motor learning. Overall, these results provide evidence that the detrimental effect of tTIS_80Hz_ on reinforcement learning of motor skills is related to modulation of neural activity specifically in the striatum.

**Figure 3.**
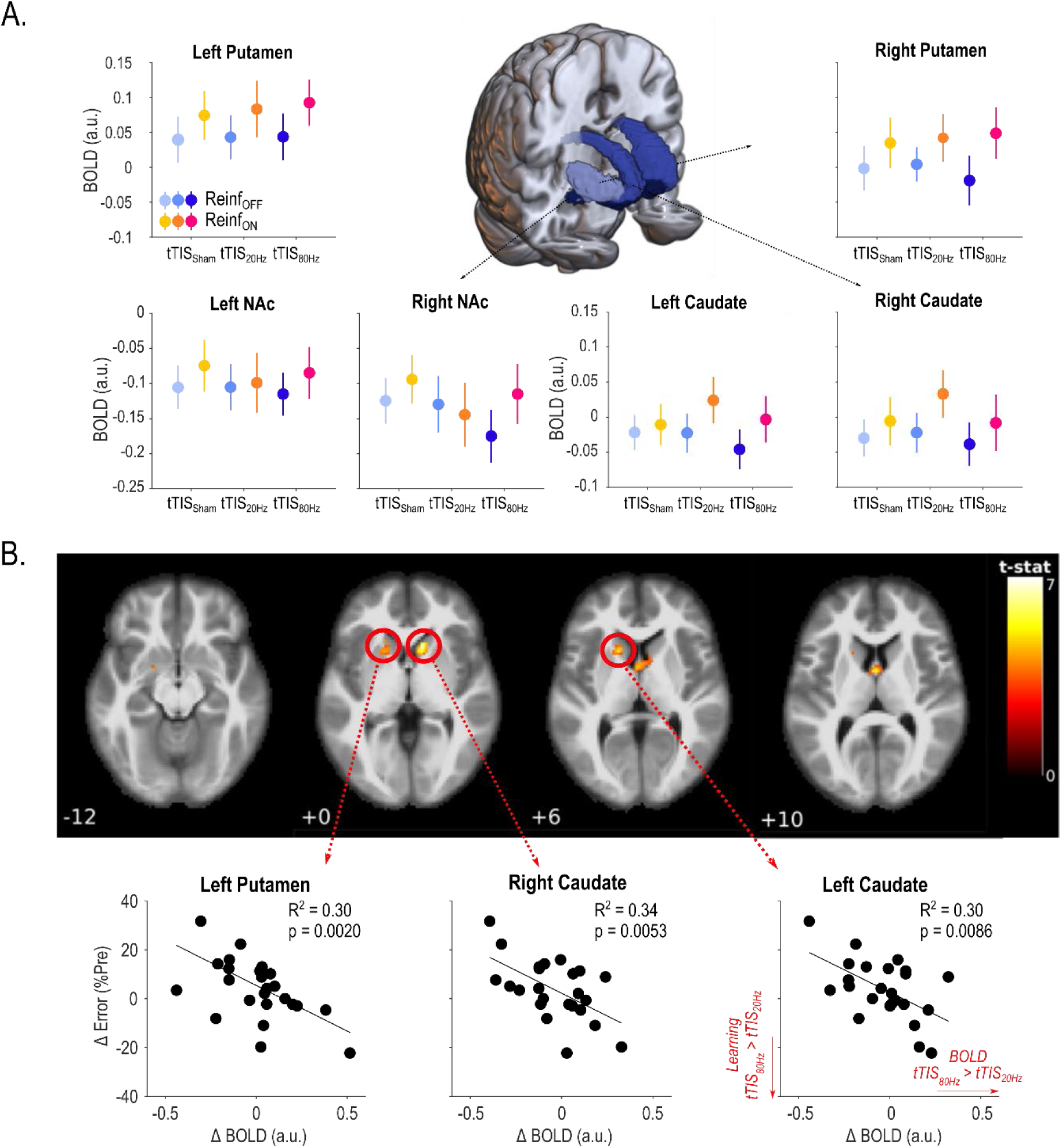
Striatal activity. **A) Striatal BOLD responses.** A 3D-reconstruction of the striatal masks used in the current experiment is surrounded by plots showing averaged BOLD activity for each mask in the different experimental conditions. A LMM ran on these data showed higher striatal responses in the Reinf_ON_ with respect to the Reinf_OFF_ condition, but no effect of tTIS_TYPE_ and no interaction. **B) Whole-brain activity associated to the behavioural effect of tTIS_80Hz_ on reinforcement motor learning.** Correlation between tTIS-related modulation of striatal activity (tTIS_80Hz_ – tTIS_20Hz_) and learning abilities in the Reinf_ON_ condition. Significant clusters of correlation were found in the left putamen and bilateral caudate (uncorrected voxel-wise FWE: p=0.001, and corrected cluster-based FDR: p=0.05). Lower panel shows individual correlations for the three significant regions highlighted in the whole-brain analysis. *: p<0.05. Data are represented as mean ± SE.

### tTIS_80Hz_ enhances effective connectivity between the striatum and frontal cortex

Interactions between the striatum and frontal cortex are crucial for a variety of behaviours including motor and reinforcement learning^13^. In particular, reinforcement motor learning requires to use information about task success to guide future motor commands^4^, a process for which the striatum may play an integrative role at the interface between fronto-striatal loops involved in reward processing and motor control^13,65^. In a subsequent analysis, we asked whether striatal tTIS modulates striatum to frontal cortex communication during reinforcement motor learning. More specifically, we computed effective connectivity (using the generalized psychophysiological interactions method^66^) between striatal and frontal regions classically associated with motor and reward-related functions, and thought to be involved in reinforcement motor learning^67,68^. For the motor network, we evaluated effective connectivity between motor parts of the striatum (i.e., dorso-lateral putamen (dlPu) and dorsal caudate (dCa)) and two regions strongly implicated in motor learning: the medial part of the supplementary motor area (SMA) and the part of the primary motor cortex (M1) associated to upper limb functions (**Figure 4A**). For the reward network, we assessed connectivity between parts of the striatum classically associated to limbic functions (i.e., the NAc and the ventro-medial putamen (vmPu) and two frontal areas involved in reward processing: the anterior cingulate cortex (ACC) and the ventro-medial prefrontal cortex (vmPFC; **Figure 4B**; ^11^). The LMM ran with the fixed effects Reinf_TYPE_, tTIS_TYPE_ and Network_TYPE_ showed a significant effect of tTIS_TYPE_ (F_(2, 2264.0)_=5.42; p=0.0045), that was due to higher connectivity in the tTIS_80Hz_ condition with respect to tTIS_Sham_ (p=0.0038; d=0.16) and tTIS_20Hz_ (at the trend level, p=0.069; d=0.11). There was no difference in connectivity between tTIS_20Hz_ and tTIS_Sham_ (p=0.58; d=0.051). Hence, tTIS_80Hz_, but not tTIS_20Hz_, enhanced effective connectivity between the striatum and frontal cortex during motor training. This increase in effective connectivity with tTIS_80Hz_ actually led to a connectivity closer to the resting state (values closer to 0, see Methods). Put differently, while the task induced a reduction in effective connectivity between striatum and frontal cortex, tTIS_80Hz_ disrupted this modulation by bringing connectivity back to the resting state.

**Figure 4.**
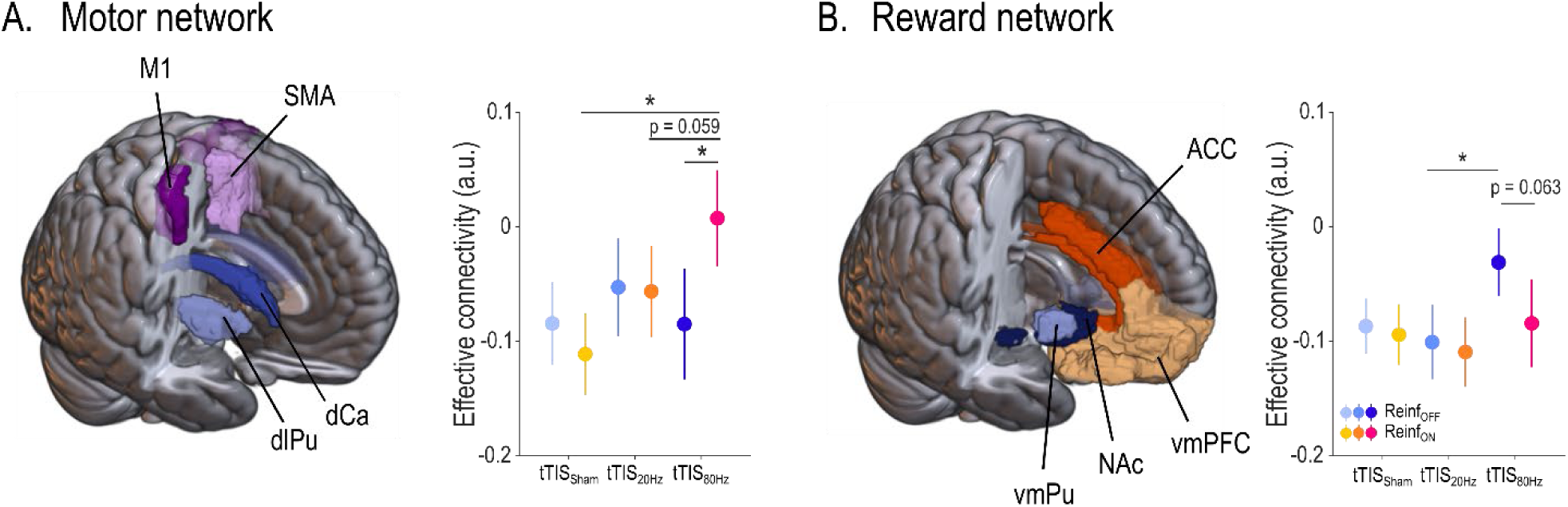
Striatum to frontal cortex effective connectivity. **A) Motor network.** On the left, 3D reconstruction of the masks used for the motor network (i.e., dorso-lateral putamen, dorsal caudate, M1, SMA). On the right, plot showing effective connectivity from motor striatum to motor cortex in the different experimental conditions. Note the increase of connectivity with tTIS_80Hz_ in the presence of reinforcement. **B) Reward network.** On the left, 3D reconstruction of the masks used for the reward network (i.e., ventro-medial putamen, NAc, vmPFC, ACC). On the right, plot showing effective connectivity from motor striatum to motor cortex in the different experimental conditions. ROIs were defined based on the BNA atlas^12^ *: p<0.05. Data are represented as mean ± SE.

The LMM did not reveal any effect of Reinf_TYPE_ (F_(1, 2264.0)_=0.010; p=0.92), Network_TYPE_ (F_(1, 2264.0)_=3.16; p=0.076) and no double interaction (note the trend for a Reinf_TYPE_ x Network_TYPE_ effect though: F_(1, 2264.0)_=3.52; p=0.061). Yet, we did find a significant Reinf_TYPE_ x tTIS_TYPE_ x Network_TYPE_ interaction (F_(2, 2264.0)_=4.87; p=0.0078). Such triple interaction was related to the fact that tTIS_80Hz_ increased connectivity in the Reinf_ON_ condition in the motor network (Reinf_ON_ vs. Reinf_OFF_: p=0.0012; d=0.33; Figure 4A), while it tended to have the opposite effect in the reward network (p=0.063; d=-0.19; Figure 4B). This increase was not present in any of the two networks when either tTIS_Sham_ or tTIS_20Hz_ were applied (all p> 0.40). Moreover, in the motor network, connectivity in the Reinf_ON_ condition was higher with tTIS_80Hz_ than with tTIS_Sham_ (p<0.001; d=0.42) and tTIS_20Hz_ (at the trend level; p=0.059; d=0.23, Figure 4A). These data suggest that tTIS_80Hz_ enhanced the neuromodulatory influence of the striatum on motor cortex during task performance, but only in the presence of reinforcement. In the reward network, post-hocs revealed that connectivity in the Reinf_OFF_ condition was significantly higher with tTIS_80Hz_ compared to tTIS_20Hz_ (p=0.045; d=0.25; Figure 4B), in line with the general effect of tTIS_TYPE_ on connectivity reported above. This pattern of results suggests that the increase of connectivity from striatum to frontal cortex observed with tTIS_80Hz_ depends on the presence of reinforcement, in particular in the motor network. Such reinforcement-dependent increase of connectivity may reflect the preferential effect of tTIS_80Hz_ on striatal gamma oscillations^69^ in a situation where these oscillations are already boosted by the presence of reinforcement^19^ (see Discussion).

In a subsequent analysis, we verified that these results did not depend on the specific frontal ROIs considered in the analysis (ROI_TYPE_: M1 and SMA in the motor network and ACC and vmPFC in the reward network). Importantly, we did not find a tTIS_TYPE_ x Reinf_TYPE_ x ROI_TYPE_ interaction neither in the motor (F_(2,1112)_=0.83; p=0.44) nor in the reward network (F_(2,1112)_=0.61; p=0.54), suggesting that the main connectivity results were consistent within a network and were not influenced by the specific frontal ROI included in the analysis (see Supplementary materials for more details on this analysis). As an additional control, we verified that the effects of tTIS_TYPE_ on connectivity could not be observed in a control network associated to language (as defined by ^70^), which was unlikely to be involved in the present task and did not include the striatum (see Methods). As expected, effective connectivity within the language network was not modulated by Reinf_TYPE_ (F_(1, 547)_=0.81; p=0.37), nor by tTIS_TYPE_ (F_(2, 547)_=0.58; p=0.56), or by Reinf_TYPE_ x tTIS_TYPE_ (F_(2, 547)_=0.45; p=0.64). Hence, tTIS and reinforcement-related changes in connectivity were consistent within the considered fronto-striatal networks and not observed in a control network unrelated to the task.

Notably, contrary to the BOLD results presented above, we did not find any correlations between the effects of tTIS_80Hz_ on connectivity and motor learning, neither in the motor (robust linear regression: tTIS_80Hz_ – tTIS_Sham_: R^2^=0.019; p=0.48; tTIS_80Hz_ – tTIS_20Hz_: R^2^=0.034; p=0.54) nor in the reward (tTIS_80Hz_ – tTIS_Sham_: R^2^=0.037; p=0.46; tTIS_80Hz_ – tTIS_20Hz_: R^2^<0.001; p=0.75) network, suggesting some degree of independence between the effect of tTIS_80Hz_ on reinforcement motor learning and on effective connectivity.

Overall, these results highlight the ability of tTIS_80Hz_, but not tTIS_20Hz_, to modulate striatum to frontal cortex connectivity, depending on the presence of reinforcement. However, the absence of correlation with the behaviour suggests that this effect may not be directly associated to the detrimental effect of tTIS_80Hz_ on reinforcement motor learning or that tTIS_80Hz_-related changes in striato-frontal communication were linked to other aspects of reinforcement learning not captured by our task.

### Neural effects of tTIS_80Hz_ depend on impulsivity

Determining individual factors that shape responsiveness to non-invasive brain stimulation approaches is a crucial step to better understand the mechanisms of action but also to envision stratification of patients in future clinical interventions^71^. A potential factor that could explain inter-individual differences in responsiveness to tTIS_80Hz_ is the level of impulsivity. As such, impulsivity has been associated to changes of gamma oscillatory activity in the striatum of rats^72^ and to the activity of fast-spiking interneurons in the striatum^73,74^, a neuronal population that is strongly entrained to gamma rythms^19,21^ and may therefore be particularly sensitive to tTIS_80Hz_. In a subsequent exploratory analysis, we asked if the neural effects of tTIS_80Hz_ were associated to impulsivity levels, as evaluated by a well-established independent delay-discounting questionnaire performed at the beginning of the experiment^75,76^. Strikingly, a whole-brain analysis revealed that impulsivity was associated to the effect of tTIS_80Hz_ on BOLD activity (with respect to tTIS_20Hz_) specifically in the left caudate nucleus (**Figure S7A, S7B, Table S4**). Moreover, the effect of tTIS_80Hz_ on striatum to motor cortex connectivity reported above was negatively correlated to impulsivity both when contrasting tTIS_80Hz_ with tTIS_Sham_ (**Figure S7C, left**) and with tTIS_20Hz_ (**Figure S7C, middle**). Such correlations were absent when contrasting tTIS_20Hz_ with tTIS_Sham_ (**Figure S7C, right**), as well as when considering the same contrasts in the reward instead of the motor network (see Supplementary materials for more details). Taken together, these results suggest that inter-individual variability in impulsivity might influence the neural responses to striatal tTIS_80Hz_.

## 3. Discussion

In this study, we combined striatal tTIS with electric field modelling, behavioural and fMRI analyses to evaluate the causal role of the striatum in reinforcement learning of motor skills in healthy humans. tTIS_80Hz_, but not tTIS_20Hz_, disrupted the ability to learn from reinforcement feedback. This behavioural effect was associated to modulation of neural activity specifically in the striatum. As a second step, we show that tTIS_80Hz_, but not tTIS_20Hz_, increased the neuromodulatory influence of the striatum on connected frontal cortical areas involved in reinforcement motor learning. Finally, inter-individual variability in the neural effects of tTIS_80Hz_ could be partially explained by impulsivity, suggesting that this trait may constitute a determinant of responsiveness to high gamma striatal tTIS. Overall, the present study shows for the first time that striatal tTIS can non-invasively modulate a striatal mechanism involved in reinforcement learning, opening new horizons for the study of causal relationships between deep brain structures and human behaviour.

We investigated the causal role of the human striatum in reinforcement learning of motor skills in healthy humans; a question that cannot be addressed with conventional non-invasive brain stimulation techniques. In particular, by stimulating at different frequencies, we aimed at dissociating striatal mechanisms involved in reinforcement and sensorimotor learning. In line with our main hypothesis, we found that striatal tTIS_80Hz_ altered reinforcement learning of a motor skill. Such disruption was frequency- and reinforcement-specific: learning was not altered with striatal tTIS_20Hz_ in the presence of reinforcement, or when striatal tTIS_80Hz_ was delivered in the absence of reinforcement. The rationale to stimulate at high gamma frequency was based on previous work showing reinforcement-related modulation of gamma oscillations in the striatum^19–21,24,26,72,77^ and in the frontal cortex^77–80^. Several neuronal mechanisms may contribute to the detrimental effect of tTIS_80Hz_ on reinforcement motor learning. First, as tTIS_80Hz_ consisted in a constant high gamma oscillating field applied on the striatum, it may have perturbed the encoding of reinforcement information into high gamma oscillations^19–21,25–27^, preventing participants to learn the motor skill based on different outcomes. Put differently, tTIS_80Hz_ may specifically saturate high gamma activity in the striatum preventing reinforcement-related modulations^81^. Moreover, because reinforcement motor learning likely engages synchronised activity in a network of regions including fronto-striatal loops, neuromodulation of a single node of the circuit may alter synchronisation of activity in the network^81^ and the temporal coordination with interacting rhythms^25^. Finally, because we did not have access to electrophysiological recordings of oscillatory activity in the striatum, the applied stimulation was not personalised as it did not take into account the individual high gamma frequency peak associated to reward processing and the potential heterogeneity of gamma activity within the striatum^24^. Hence, tTIS_80Hz_ may have resulted in a frequency mismatch between the endogenous high gamma activity and the externally imposed rhythm, that could paradoxically result in a reduction of neuronal entrainment, in particular when the frequency mismatch is relatively low^82^. Importantly, in contrast to striatal tTIS_80Hz_, we found that tTIS_20Hz_ reduced learning, but only in the absence of reinforcement. This result fits well with the literature linking striatal beta oscillations to sensorimotor functions^28,29,31,83–85^. Taken together, an interpretation of these results is that different oscillations within the striatum support qualitatively distinct motor learning mechanisms with beta activity contributing mostly to sensory-based learning and high gamma activity being particularly important for reinforcement learning. This being said, it is important to note that because we do not have concurrent electrophysiological recordings within the striatum, we cannot be sure that the effects of tTIS_20Hz_ and tTIS_80Hz_ were related to frequency-specific interactions with beta or high gamma rhythms respectively, or rather resulted from different broadband responses when stimulating at these frequencies. Yet, these results still suggest that sensory- and reinforcement-based motor learning rely on partially different neural mechanisms, in line with previous literature^8,9,61,68,86,87^.

Interestingly, striatal tTIS also impaired tracking performance during training, irrespective of the presence of reinforcement. This frequency-dependent reduction of motor performance may be due to altered neuronal processing in the sensorimotor striatum that may lead to less fine-tuned motor control abilities^88^. Importantly though, tTIS did not modulate the ability of participants to benefit from real-time reinforcement feedback during motor performance. This suggests that striatal tTIS_80Hz_ altered the beneficial effects of reinforcement on learning (as evaluated in Test conditions at Post-training), but not on motor performance (as evaluated during Training). Such dissociation between the effects of striatal tTIS_80Hz_ on reinforcement-related gains in motor performance and learning may be explained by the fact that these two phases of the protocol probe different processes^7,54,56,89–91^. While improvement of motor performance with reinforcement relies on rapid feedback corrections based on expected outcomes^67,92–95^, reinforcement gains in learning (i.e., probed in Test conditions without reinforcement) may rather reflect the beneficial effect of reinforcement on the retention of motor memories^5,7,54,90^. This idea that mechanisms underlying performance changes in training and retention phases are partially different is well supported by previous motor learning literature^6,8,96^. For instance, in sensorimotor adaptation paradigms, the presence of reward boosts motor memory retention but not the adaptation process itself^7,97^, and M1 transcranial direct current stimulation modulates the effect of reward on retention but has no effect on the training phase^90^. Hence, a potential explanation for the present results is that striatal tTIS_80Hz_ did not disrupt rapid motor corrections based on recent outcomes during training, but may rather alter the strengthening of the memory trace based on reinforcements^6,7^. Overall, these results are compatible with the view that specific patterns of oscillatory activity in the striatum are involved in motor control and learning processes^31^, and can be modulated with electrical stimulation^69,98,99^.

To better understand the neural effects and frequency-specificity of tTIS, we coupled striatal tTIS and task performance with simultaneous fMRI acquisition. The imaging results support the view that the effect of tTIS_80Hz_ on reinforcement learning of motor skills was indeed related to neuromodulation of the striatum. As such, when considering averaged BOLD activity, we found a general increase of striatal activity when reinforcement was provided^11^, but no effect of tTIS. Crucially though, the detrimental effect of tTIS_80Hz_ on reinforcement learning was related to a specific modulation of activity in the caudate and putamen, providing evidence that the present behavioural effects were indeed driven by focal neuromodulation of the striatum (Figure 3). Interestingly, participants with stronger disruption of reinforcement learning at the behavioural level were also the ones exhibiting stronger suppression of striatal activity with tTIS_80Hz_ (compared to tTIS_20Hz_), suggesting that tTIS-induced reduction of striatal activity is detrimental for reinforcement motor learning. Further analyses showed that tTIS_80Hz_, but not tTIS_20Hz_, increased the neuromodulatory influence of the striatum on frontal areas known to be important for motor learning and reinforcement processing^96,100^. More specifically, tTIS_80Hz_ disrupted the task-related decrease in connectivity observed with tTIS_Sham_ and tTIS_20Hz_, bringing connectivity closer to resting-state values. Interestingly, this effect depended on the type of network considered (reward vs. motor) and on the presence of reinforcement. Striatal tTIS_80Hz_ coupled with reinforcement increased connectivity between the motor striatum and the motor cortex while it tended to have the opposite effect when considering the connectivity between limbic parts of the striatum and pre-frontal areas involved in reward processing (Figure 4). This result may reflect the differential influence of striatal tTIS on distinct subparts of the striatum, depending on their pattern of activity during the task^52^. As such, a recent study in non-human primates showed that tACS can have opposite effects on neuronal activity based on the initial entrainment of neurons to the target frequency^82^. Hence, the present differential effects of tTIS_80Hz_ on motor and reward striato-frontal pathways may be due to different initial patterns of activity in these networks in the presence of reinforcement. Electrophysiological recordings with higher temporal resolution than fMRI are required to confirm or infirm this hypothesis. Overall, the present neuroimaging results support the idea that the behavioural effects of striatal tTIS_80Hz_ on reinforcement learning are associated to a selective modulation of striatal activity that influences striato-frontal communication.

The fact that we observed increased connectivity with tTIS_80Hz_ and at the same time a disruption of behaviour may appear contradictory at first glance. Yet, multiple lines of evidence indicate that increases in connectivity are not necessarily beneficial for behaviour. For instance, the severity of motor symptoms in Parkinson’s disease is associated with excessive connectivity in the beta band and reduction of such connectivity with treatment is associated to clinical improvement^29,32^. Moreover, there is evidence that excessive functional^101,102^ as well as structural^103,104^ connectivity in fronto-striatal circuits is associated to impulsivity. Hence, the increase in connectivity observed with tTIS_80Hz_ appears to be compatible with the behavioural findings. This being said, contrary to the BOLD results, we did not find any correlation between the effects of tTIS_80Hz_ on connectivity and on reinforcement motor learning, suggesting some degree of independence between these two effects. Future studies could aim at determining if tTIS_80Hz_-related changes in striato-frontal communication are linked to other aspects of reward processing, not captured by our reinforcement motor learning task.

From a methodological point of view, the present results provide new experimental support to the idea that the effects of tTIS are related to amplitude modulation of electric fields deep in the brain and not to the high frequency fields themselves, in line with recent work^41,42,52^. As such, the different behavioural and neural effects of striatal tTIS_80Hz_ and tTIS_20Hz_ despite comparable carrier frequencies (centered on 2kHz) indicate that temporal interference was indeed the driving force of the present effects. Moreover, disruption of reinforcement motor learning with tTIS_80Hz_ (relative to tTIS_20Hz_) was specifically related to neuromodulation of the striatum, where the amplitude of the tTIS field was highest according to our simulations (see ^51,53^ for recent validations of comparable simulations in cadavers experiments). Hence, we believe that the frequency- and reinforcement-dependent tTIS effects reported here cannot be explained by direct modulation of neural activity by the high frequency fields. Yet, disentangling the neural effects of the low-frequency envelope and the high frequency carrier appears as an important next step to better characterise the mechanisms underlying tTIS^47^. We also note that the tTIS field strengths achieved according to our simulations (in the range of 0.5-0.6 V/m) were sufficient to induce behavioral and neural effects, in line with recent data^52,53^ (see also ^48^). Determining the minimum effective dose for tTIS is an important line of future research given recent simulation results suggesting that stimulation via an amplitude modulation with high frequency carrier signals (such as arising during tTIS) may require higher dosages compared to conventional electrical stimulation with low frequencies (such as during tACS), likely due to the low-pass filtering properties of neurons^43,105^.

Finally, the strength of the behavioural effects of tTIS can be considered small to medium^106^ (d=0.2-0.5). We note that these effect sizes are consistent with studies applying other types of non-invasive brain stimulation in healthy young adults, both in the context of motor learning (see ^107^ for a meta-analysis), and reward tasks (e.g., ^108,109^), despite the much longer stimulation time used in these studies (between 3 and 20 times longer). Overall, albeit moderate, we believe that the present effect sizes are relevant and consistent with what can be expected from the non-invasive brain stimulation literature.

### Limitations

The present study includes some limitations that we would like to acknowledge. First, at the imaging level, we did not find a significant effect of reinforcement at the whole-brain level. This might be due to the short duration of the task (6×40s), combined with the fact that we did not couple reinforcement to monetary incentives, a manipulation known to strongly boost striatal activity in the context of motor learning^18^. Yet, when considering BOLD activity in the striatal ROIs, we did find a significant effect of reinforcement, suggesting that our experimental manipulation did increase striatal activity but that the strength of the effect was insufficient to survive at the whole-brain level. Second, we did not find any effect of tTIS when considering averaged BOLD activity. Again, the short duration of the blocks may contribute to this non-significant effect. Another possible interpretation is that the effect of tTIS on BOLD activity is not uniform across participants as it likely depends on individual anatomy and function of the targeted brain region, as observed for other non-invasive brain stimulation techniques^110^. Consistently, we found a correlation between levels of impulsivity and the neural effects of tTIS_80Hz_ (both BOLD and connectivity, Figure S7). Importantly though, when including learning as a behavioral regressor we did find significant clusters of correlation specifically in the striatum (Figure 3), suggesting that the behavioural effects were indeed related to modulation of activity in the target region. Notably, this result was significant when contrasting tTIS_80Hz_ to the active control (tTIS_20Hz_), but not to tTIS_Sham_. Overall, we believe that the fMRI data does provide interesting support that the behavioural effects of the stimulation were indeed related to modulation of neural activity in the striatum, also in line with the present simulations on realistic head models (Figure 1) and the connectivity results (Figure 4). This idea is also in agreement with another recent study investigating the effects of tTIS on motor sequence learning^52^. Notably though, a limitation of the present dataset is the very short duration of stimulation and imaging for each experimental condition, that may explain some inconsistencies in the results. Hence, following this first proof-of-concept study showing robust behavioural effects and related neural changes, future studies including longer fMRI and stimulation sessions are required to further confirm these results.

Finally, within the present study the computational modelling was performed on a realistic, detailed head model (i.e., the MIDA model^59^, see Methods). One limitation of this approach is that the electric field simulations do not take individual structural information into account. Such individual modeling would require information on brain anisotropy, an aspect that is likely to significantly influence tTIS exposure^44,111^. However, in the present study diffusion MRI to evaluate fractional anisotropy was not acquired. Future studies including diffusion MRI data will allow for personalised modelling, paving the way for individualised tTIS informed by brain structure^53^.

## Conclusion

The present findings show for the first time the ability of non-invasive striatal tTIS to interfere with reinforcement learning in humans through a selective modulation of striatal activity and support the causal functional role of the human striatum in reinforcement motor learning. This deep brain stimulation was well tolerated and compatible with efficient blinding, suggesting that tTIS provides the exciting option to circumvent the steep depth-focality trade-off of current non-invasive brain stimulation approaches in a safe and effective way. Overall, tTIS opens new possibilities for the study of causal brain-behaviour relationships and for the treatment of neuro-psychiatric disorders associated to alterations of deep brain structures.

## 4. Methods

### 4.1. Participants

A total of 48 right-handed healthy volunteers participated in the study. 24 participants were enrolled for the main tTIS study (15 women, 25.3 ± 0.7 years old; mean ± SE). Another group of 24 volunteers participated in the behavioural control experiment (Figure S2, 14 women, 24.2 ± 0.5 years old). Handedness was determined via a shortened version of the Edinburgh Handedness inventory^112^ (laterality index = 89.3 ± 2.14% for the main study and 86.4 ± 2.51% for the control experiment). None of the participants suffered from any neurological or psychiatric disorder, nor taking any centrally-acting medication (see Supplementary Materials for a complete list of exclusion criteria). All participants gave their written informed consent in accordance with the Declaration of Helsinki and the Cantonal Ethics Committee Vaud, Switzerland (project number 2020-00127). Finally, all participants were asked to fill out a delay-discounting monetary choice questionnaire^113^, which evaluates the propensity of subjects to choose smaller sooner rewards over larger later rewards, a preference commonly associated to choice impulsivity^75,114^.

### 4.2. Experimental procedures

The study employed a randomised, double-blind, sham-controlled design. Following screening and inclusion, participants were invited to a single experimental session including performance of a motor learning task with concurrent transcranial electric Temporal Interference stimulation (tTIS) of the striatum and functional magnetic resonance imaging (fMRI). Overall, participants practiced 6 blocks of trials, that resulted from the combination of two reinforcement feedback conditions (Reinf_TYPE_: Reinf_ON_ or Reinf_OFF_) with three types of striatal stimulation (tTIS_TYPE_: tTIS_Sham_, tTIS_20Hz_ or tTIS_80Hz_).

#### 4.2.1. Motor learning task

##### 4.2.1.1. General aspects

Participants practiced an adaptation of a widely used force-tracking motor task^54,55^ with a fMRI-compatible fiber optic grip force sensor (Current designs, Inc., Philadelphia, PA, USA) positioned in their right hand. The task was developed on Matlab 2018 (the Mathworks, Natick, Massachusetts, USA) exploiting the Psychophysics Toolbox extensions^115,116^ and was displayed on a computer screen with a refresh rate of 60 Hz. The task required participants to squeeze the force sensor to control a cursor displayed on the screen. Increasing the exerted force resulted in the cursor moving vertically and upward in a linear way. Each trial started with a preparatory period in which a sidebar appeared at the bottom of the screen (**Figure 1A**). After a variable time interval (0.9 to 1.1 s), a cursor (black circle) popped up in the sidebar and simultaneously a target (grey larger circle with a cross in the middle) appeared, indicating the start of the movement period. Subjects were asked to modulate the force applied on the transducer to keep the cursor as close as possible to the center of the target. The target moved in a sequential way along a single vertical axis for 7 s. The maximum force required (i.e., the force required to reach the target when it was in the uppermost part of the screen; MaxTarget_Force_) was set at 4% of maximum voluntary contraction (MVC) evaluated at the beginning of the experiment. This low force level was chosen based on pilot experiments to limit muscular fatigue. Finally, each trial ended with a blank screen displayed for 2 s before the beginning of the next trial.

##### 4.2.1.2. Trial types and reinforcement manipulation

During the experiment, participants were exposed to different types of trials (**Figure 1A, Video S1**). In Test trials, the cursor remained on the screen and the target was consistently displayed in grey for the whole duration of the trial. These trials served to evaluate Pre- and Post-training performance for each block, without any disturbance. In Reinf_ON_ and Reinf_OFF_ trials (used during Training only), we provided only partial visual feedback to the participants in order to increase the impact of reinforcement on learning^4,56–58^. As such, the cursor was only intermittently displayed during the trial: it was always displayed in the first second of the trial, and then disappeared for a total of 4.5 s randomly split on the remaining time by bits of 0.5 s. The cursor was therefore displayed 35.7% of the time during these trials (2.5 s over the 7 s trial). Importantly, contrary to the cursor, the target always remained on the screen for the whole trial and participants were instructed to continue to track the target even when the cursor was away.

In addition to this visual manipulation, in Reinf_ON_ trials, participants also trained with reinforcement feedback indicating success or failure of the tracking in real time. As such, participants were informed that, during these trials, the color of the target would vary as a function of their performance: the target was displayed in green when tracking was considered as successful and in red when it was considered as failure. Online success on the task was determined based on the Error, defined as the absolute force difference between the force required to be in the center of the target and the exerted force^4,54–56^. The Error, expressed in percentage of MVC, was computed for each frame refresh and allowed to classify a sample as successful or not based on a closed-loop reinforcement schedule^8^. More specifically, for each training trial, a force sample (recorded at 60 Hz, corresponding to the refresh rate of the monitor) was considered as successful if the computed Error was below the median Error over the 4 previous trials at this specific sample. Put differently, to be successful, participants had to constantly beat their previous performance. This closed-loop reinforcement schedule allowed us to deliver consistent reinforcement feedback across individuals and conditions (see control analysis on success rates in the Supplementary materials), while maximizing uncertainty on the presence of reinforcement, an aspect that is crucial for efficient reinforcement motor learning^117^. Notably, in addition to this closed-loop design, samples were also considered as successful if the cursor was very close to the center of the target (i.e., within one radius around the center, corresponding to an Error below 0.2% of MVC). This was done to prevent any conflict between visual information (provided by the position of the cursor relative to the target) and reinforcement feedback (provided by the color of the target), which could occur in situations of extremely good performance (when the closed-loop Error cut-off is below 0.2% of MVC).

As a control, Reinf_OFF_ trials were similar to Reinf_ON_ trials with the only difference that the displayed colors were either cyan or magenta, and were generated randomly. Participants were explicitly told that, in this condition, colors were displayed randomly and could be ignored. The visual properties of the target in the Reinf_OFF_ condition were designed to match the Reinf_ON_ condition in terms of relative luminance (cyan: RGB = [127.5 242.1 255] matched to green: [127.5 255 127.5] and magenta: [211.7 127.5 255] to red: [255 127.5 127.5]) and average frequency of change in colors (i.e., the average number of changes in colors divided by the total duration of a trial, see Supplementary materials).

Notably, in this task, Training trials differed from Test trials regarding not only the color of the target (red/green or cyan/magenta in Training trials and grey in Test trials) but also the visual feedback experienced (partial and full visual feedback in Training and Test trials, respectively). This choice was motivated by several reasons. First, we wanted to evaluate learning in the classical, unperturbed, version of the force-tracking task^54,55^, which is compatible with clinical translation. Second, based on additional behavioural data on another group of participants (n = 24, see Figure S2), we found that significant effects of reinforcement on learning were observed only when training was performed with partial visual feedback (displayed on 35.7% of the trial time, as in the present study), in line with previous results^57,62^. However, this additional study also revealed very limited improvement of performance during training with partial visual feedback, potentially due to ceiling effects on performance in this condition. Yet, the improvement of performance when comparing the Pre and Post-training assessments strongly suggested that practicing the task with partial visual feedback still induced significant learning of the skill. Finally, the change in visual feedback between Training and Post-training was the same in all experimental conditions; this aspect of the task is therefore unlikely to explain the reinforcement as well as the stimulation effects reported here.

Even though our study focused on reinforcement motor learning, it is worth mentioning that other learning mechanisms such as error-based or strategic processes are likely to be also engaged during the force-tracking task and may have recruited other brain regions beyond the striatum (Spampinato and Celnik, 2020). Notably though, our protocol was specifically designed to compare in the same individuals, learning in Reinf_ON_ and Reinf_OFF_ conditions while keeping the other parameters of the task constant, to specifically isolate the contribution of reinforcement processes in motor learning.

##### 4.2.1.3. Motor learning protocol

After receiving standardised instructions about the force-tracking task, participants practiced 5 blocks of familiarization (total of 75 trials) without tTIS. The first block of familiarization included 20 trials with the target moving in a regular fashion (0.5 Hz sinuoid). Then, in a second block of familiarization, participants performed 35 trials of practice with an irregular pattern, with the same properties as the training patterns (see below). Finally, we introduced the reinforcement manipulation and let participants perform 2 short blocks (8 trials each) including Reinf_ON_ and Reinf_OFF_ trials. These four first blocks of familiarization were performed outside the MRI environment. A last familiarization block (4 trials) was performed after installation in the scanner, to allow participants to get used to performing the task in the MRI. This long familiarization allowed participants to get acquainted with the use of the force sensor, before the beginning of the experiment.

During the main part of the experiment, participants performed 6 blocks of trials in the MRI with concurrent striatal tTIS (**Figure 1B**). Each block was composed of 4 Pre-training trials followed by 24 Training and 8 Post-training trials. Pre- and Post-training trials were performed in Test conditions, without tTIS and were used to evaluate motor learning. Training trials were performed with or without reinforcement feedback and with concomitant striatal tTIS and were used as a proxy of motor performance. During Training, trials were interspersed with 25 s resting periods every 4 trials (used for fMRI contrasts, see below). The order of the 6 experimental conditions was pseudo-randomised across participants: the 6 blocks were divided into 3 pairs of blocks with the same tTIS condition and each pair was then composed of one Reinf_ON_ and one Reinf_OFF_ block. Within this structure, the order of the tTIS_TYPE_ and Reinf_TYPE_ conditions were balanced among the 24 participants. Hence, this randomisation allowed us to ensure that any order effect that may arise from the repetition of the learning blocks would have the same impact on each experimental condition (e.g., 4 subjects experienced tTIS_80Hz_ -Reinf_ON_ in the first block, 4 other subjects in the second block, 4 in the third block etc.).

As mentioned above, the protocol involved multiple evaluations of motor learning within the same experimental session. In order to limit carry-over effects from one block to the following, each experimental block was associated to a different pattern of movement of the target (**Figure S1**). Put differently, in each block, participants had to generate a new pattern of force to successfully track the target. To balance the patterns’ difficulty, they all consisted in the summation of 5 sinusoids of variable frequency (range: 0.1-1.5 Hz) that presented the following properties: a) Average force comprised between 45 and 55% of the MaxTarget_Force_; b) Absolute average derivative comprised between 54 and 66 % of the MaxTarget_Force_/s; c) Number of peaks = 14 (defined as an absolute change of force of at least 1% of MaxTarget_Force_). These parameters were determined based on pilot experiments to obtain a relevant level of difficulty for young healthy adults and consistent learning across the different patterns.

#### 4.2.2. Transcranial Electric Temporal Interference Stimulation (tTIS) applied to the striatum

##### 4.2.2.1. General concept

Transcranial temporal interference stimulation (tTIS) is an innovative non-invasive brain stimulation approach, in which two or more independent stimulation channels deliver high-frequency currents in the kHz range (oscillating at f1 and f1 + Δf; **Figure 1C**). These high-frequency currents are assumed to be too high to effectively modulate neuronal activity ^41,50,118^. Still, by applying a small shift in frequency, they result in a modulated electric field with the envelope oscillating at the low-frequency Δf (target frequency) where the two currents overlap. The peak of the modulated envelope amplitude can be steered towards specific areas located deep in the brain, by tuning the position of the electrodes and current ratio across stimulation channels^41^ (**Figure 1C, 1D**). Based on these properties, tTIS has been shown to be able to focally target activity of deep structures in rodents, without engaging overlying tissues^41^. Here, we applied temporal interference stimulation transcranially via surface electrodes applying a low-intensity, sub-threshold protocol following the currently accepted cut-offs and safety guidelines for low-intensity transcranial electric stimulation in humans^119^.

##### 4.2.2.2. Stimulators

The currents for tTIS were delivered by two independent DS5 isolated bipolar constant current stimulators *(Digitimer Ltd, Welwyn Garden City, UK)*. The stimulation patterns were generated using a custom-based Matlab graphical user interface and transmitted to the current sources using a standard digital-analog converter *(DAQ USB-6216, National Instruments, Austin, TX, USA)*. Finally, an audio transformer was added between stimulators and subjects, in order to avoid possible direct current accumulation.

##### 4.2.2.3. Stimulation protocols

During the 6 Training blocks, we applied three different types of striatal tTIS (2 blocks each): a stimulation with a tTIS envelope modulated at 20Hz (tTIS_20Hz_), a stimulation with a tTIS envelope modulated at 80Hz (tTIS_80Hz_) and a sham stimulation (tTIS_Sham_). For tTIS_20Hz_, the posterior stimulation channel (TP7-TP8, see below) delivered a 1.99 kHz stimulation while the anterior one delivered a 2.01 kHz (Δf = 20 Hz). For tTIS_80Hz_, the posterior and anterior channels delivered 1.96 kHz and 2.04 kHz, respectively (Δf = 80 Hz). Hence in both conditions, the high frequency component was comparable and the only difference was Δf. During each block, tTIS was applied for 5 minutes (6 x 50 s) during Training. Each stimulation period started and ended with currents ramping-up and -down, respectively, for 5 s. tTIS was applied only while participants were performing the motor task and not during resting periods or Pre- and Post-training assessments. Finally, tTIS_Sham_ consisted in a ramping-up (5 s) immediately followed by a ramping-down (5 s) of 2 kHz currents delivered without any shift in frequency. This condition allowed us to mimic the sensations experienced during the active conditions tTIS_20Hz_ and tTIS_80Hz_, while delivering minimal brain stimulation (Figure S5). A trigger was sent 5 seconds before the beginning of each trial in order to align the beginning of the task and the beginning of the frequency shift after the ramp-up. Other tTIS parameters were set as follows: current intensity per stimulation channel = 2 mA (baseline-to-peak), electrode type: round, conductive rubber with conductive cream/paste, electrode size = 3 cm^2^ (see ContES checklist in Supplementary materials for more details).

The stimulation was applied within the MRI environment (Siemens 3T MAGNETOM Prisma; Siemens Healthcare, Erlangen, Germany) using a standard RF filter module and MRI-compatible cables *(neuroConn GmbH, Ilmenau, Germany)*. The technological, safety and noise tests, and methodological factors can be found in Supplementary materials (Table S1) and are based on the ContES Checklist ^120^.

##### 4.2.2.4. Modelling

Electromagnetic simulations were carried out to identify optimised electrode placement and current steering parameters. Simulations were performed using the MIDA head model^59^, a detailed anatomical head model featuring >100 distinguished tissues and regions that was derived from multi-modal image data of a healthy female volunteer. Importantly, for brain stimulation modelling, the model differentiates different scalp layers, skull layers, grey and white matter, cerebrospinal fluid, and the dura and accounts for electrical conductivity anisotropy and neural orientation based on diffusion tensor imaging (DTI) data. Circular electrodes (radius = 0.7 cm) were positioned on the skin according to the 10-10 system and the electromagnetic exposure was computed using the ohmic-current-dominated electro-quasistatic solver from Sim4Life v5.0 (ZMT Zurich MedTech AG, Switzerland), which is suitable due to the dominance of ohmic currents over displacement currents and the long wavelength compared with the simulation domain^121^. Dielectric properties were assigned based on the IT’IS Tissue Properties Database v4.0^122^. Rectilinear discretization was performed, and grid convergence as well as solver convergence analyses were used to ensure negligible numerical uncertainty, resulting in a grid that included more than 54M voxels. Dirichlet voltage boundary conditions, and then current normalization were applied. The electrode-head interface contact was treated as ideal. tTIS exposure was quantified according to the maximum modulation envelope magnitude formula from Grossman et al., (2017)^41^. Then, a sweep over 960 permutations of the four electrode positions was performed, considering symmetric and asymmetric montages with parallel (sagittal and coronal) or crossing current paths, while quantifying bilateral striatum (putamen [BNA regions 225, 226, 229, 230], caudate [BNA regions 219, 220, 227, 228] and nucleus accumbens [BNA regions 223, 224]) exposure performance according to three metrics: a) target exposure strength, b) focality ratio (the ratio of target tissue volume above threshold compared to the whole-brain tissue volume above threshold, a measure of stimulation selectivity), and c) activation ratio (percentage of target volume above threshold with respect to the total target volume, a measure of target coverage). We defined the threshold as the 98^th^ volumetric iso-percentile level of the tTIS. From the resulting Pareto-optimal front, two configurations stood out particularly: one that maximised focality and activation (AF3 - AF4, P7 - P8) and a second one that accepts a reduction of these two metrics by a quarter, while increasing the target exposure strength by more than 50% (F3-F4, TP7-TP8). This last montage was selected, to ensure sufficient tTIS exposure in the striatum^52^ (Figure 1C, 1D).

##### 4.2.2.5. Electrode positioning and evaluation of stimulation-associated sensations

Based on the modelling approach described above, we defined the stimulation electrode positions in the framework of the EEG 10-10 system^123^. The optimal montage leading in terms of target (i.e. the bilateral striatum) exposure strength and selectivity, was composed of the following electrodes: F3, F4, TP7 and TP8. Their locations were marked with a pen on the scalp and, after skin preparation (cleaned with alcohol), round conductive rubber electrodes of 3 cm^2^ were placed adding a conductive paste (*Ten20, Weaver and Company, Aurora, CO, USA or Abralyt HiCl, Easycap GmbH, Woerthsee-Etterschlag, Germany*) as an interface to the skin. Electrodes were held in position with tape and cables were oriented towards the top in order to allow good positioning inside the scanner. Impedances were checked and optimised until they were below 20 kΩ ^48^. Once good contact was obtained, we tested different intensities of stimulation for each stimulation protocol in order to familiarise the participants with the perceived sensations and to systematically document them. tTIS _Sham_, tTIS_20Hz_ and tTIS_80Hz_ were applied for 20 seconds with the following increasing current amplitudes per channel: 0.5 mA, 1 mA, 1.5 mA and 2 mA. Participants were asked to report any kind of sensation and, if a sensation was felt, they were asked to grade the intensity from 1 to 3 (light to strong) as well as give at least one adjective to describe it (Figure S5). Following this step, cables were removed to be replaced by MRI-compatible cables and a bandage was added to apply pressure on the electrodes and keep them in place. An impedance check was repeated in the MRI right before the training and then again at the end of all recordings.

#### 4.2.3. MRI data acquisition

Structural and functional images were acquired using a 3T MAGNETOM PRISMA scanner (*Siemens, Erlangen, Germany*). T1-weighted images were acquired via the 3D MPRAGE sequence with the following parameters: TR = 2.3 s; TE = 2.96 ms; flip angle = 9°; slices = 192; voxel size = 1 × 1 × 1 mm, FOV = 256 mm. Anatomical T2 images were also acquired with the following parameters: TR = 3 s; TE = 409 ms; flip angle = 120°; slices = 208; voxel size = 0.8 × 0.8 × 0.8 mm, FOV = 320 mm. Finally, functional images were recorded using Echo-Planar Imaging (EPI) sequences with the following parameters: TR = 1.25 s; TE = 32 ms; flip angle = 58°; slices = 75; voxel size = 2 × 2 × 2 mm; FOV = 112 mm.

### 4.3. Data and statistical analyses

Data and statistical analyses were carried out with Matlab 2018a (the Mathworks, Natick, Massachusetts, USA) and the R software environment for statistical computing and graphics (R Core Team 2021, Vienna, Austria). Robust linear regressions were fitted with the Matlab function robustfit. Linear mixed models (LMM) were fitted using the lmer function of the lme4 package in R^124^. As random effects, we added intercepts for participants and block. Normality of residuals, and homoscedasticity of the data were systematically checked, and logarithmic transformations were applied when necessary (i.e., when skewness of the residuals’ distribution was not comprised between - 2 and 2^125^ or when homoscedasticity was violated based on visual inspection). To mitigate the impact of isolated influential data points on the outcome of the final model, we used tools of the influence.ME package to detect and remove influential cases based on the following criterion: distance > 4 * mean distance^126^. Statistical significance was determined using the anova function with Satterthwaite’s approximations of the lmerTest package^127^. For specific post-hoc comparisons we conducted pairwise comparisons by computing estimated marginal means with the emmeans package with Tukey adjustment of p-values^128^. Standardised effect size measures were obtained using the eff_size function of the emmeans package^129^. The level of significance was set at p<0.05.

#### 4.3.1. Behavioural data

##### 4.3.1.1. Evaluation of motor learning

The main goal of the present study was to evaluate the influence of striatal tTIS on reinforcement motor learning. To do so, we first removed trials, in which participants did not react within 1 s after the appearance of the cursor and target, considering that these extremely long preparation times may reflect significant fluctuations in attention^130^. This occurred extremely rarely (0.52 % of the whole data set). For each subject and each trial, we then quantified the tracking Error as the absolute force difference between the applied and required force as done previously^4,54,56^. Tracking performance during Training and Post-training trials were then normalised according to subjects’ initial level by expressing the Error data in percentage of the average Pre-training Error for each block. In order to test our main hypothesis predicting specific effects of striatal tTIS on reinforcement motor learning, we performed a LMM on the Post-training data with tTIS_TYPE_ and Reinf_TYPE_ as fixed effects. We then also ran the same analysis on the Training data, to evaluate if striatal tTIS also impacted on motor performance, while stimulation was being delivered.

As a control, we checked that initial performance at Pre-training was not different between conditions with a LMM on the Error data obtained at Pre-training. Again, tTIS_TYPE_ and Reinf_TYPE_ were considered as fixed effects. Finally, another LMM was fitted with the fixed effect tTIS_TYPE_ to verify that the amount of positive reinforcement (as indicated by a green target) in the Reinf_ON_ blocks was similar across tTIS_TYPES_.

#### 4.3.2. fMRI data

##### 4.3.2.1. Imaging Preprocessing

We analyzed functional imaging data using Statistical Parametric Mapping 12 (*SPM12; The Wellcome Department of Cognitive Neurology, London, UK*) implemented in MATLAB R2018a (*Mathworks, Sherborn, MA*). All functional images underwent a common preprocessing including the following steps: slice time correction, spatial realignment to the first image, normalization to the standard MNI space and smoothing with a 6 mm full-width half-maximal Gaussian kernel. T1 anatomical images were then co-registered to the mean functional image and segmented. This allowed to obtain bias-corrected gray and white matter images, by normalizing the functional images via the forward deformation field. To select subjects with acceptable level of head movement, framewise displacement was calculated for each run. A visual check of both non-normalised and normalised images was performed in order to ensure good preprocessing quality. Finally, possible tTIS-related artifacts were investigated based on signal to noise ratio maps (see below).

##### 4.3.2.2. Signal to Noise Ratio

Total signal to noise ratio (tSNR) maps were computed to check the presence of possible artifacts induced by the electrical stimulation. The values were calculated per each voxel by dividing the mean of the voxel time series by its standard deviation. Spherical regions of interest were then defined both underneath the tTIS electrodes and at 4 different locations, distant from the electrodes as a control. The center of each spherical ROI was obtained by projecting the standard MNI coordinates of each electrode on the scalp^131^ toward the center of the brain. After visual inspection of the ROIs, average tSNR maps were extracted within each sphere. A LMM was used to compare the average SNR underneath the electrodes versus the control regions and between stimulation protocols. The results of this analysis are presented in Supplementary materials (Figure S8).

##### 4.3.2.3. Task-based BOLD activity analysis

A general linear model was implemented at the single-subject level in order to estimate signal amplitude. Eight regressors were included in the model: 6 head motion parameters (displacement and rotation) and normalised time series within the white matter and the corticospinal fluid. Linear contrasts were then computed to estimate specific activity during the motor task with respect to resting periods. Functional activation was also extracted within specific ROIs individually defined based on structural images. More specifically, the Freesurfer recon-all function was run based on the structural T1w and T2w images (https://surfer.nmr.mgh.harvard.edu/). The BNA parcellation was derived on the individual subject space and the selected ROIs were then co-registered to the functional images and normalised to the MNI space. BOLD activity within the individual striatal masks was averaged and compared between different striatal nuclei namely the putamen (BNA regions 225, 226, 229, 230), caudate (BNA regions 219, 220, 227, 228) and nucleus accumbens (BNA regions 223, 224). Comparison between conditions were presented for uncorrected voxel-wise FWE, p=0.001 and multiple comparison corrected at the cluster level to reduce False Discovery Rate (FDR), p=0.05.

##### 4.3.2.4. Effective connectivity analyses

As an additional investigation, we computed task-modulated effective functional connectivity by means of the CONN toolbox 2021a (www.nitrc.org/projects/conn, RRID:SCR_009550) running in Matlab R2018a (*Mathworks, Sherborn, MA*). An additional denoising step was added by applying a band-pass filtering from 0.01 to 0.1 Hz and by regressing potential confounders (white matter, CSF and realignment parameters). After that, generalized Psycho-Physiological Interactions (gPPI) connectivity was extracted within specific pre-defined customised sub-networks: a reward and a motor network. gPPI evaluates condition-specific changes in effective connectivity, defined as the directed effect that one brain region has on another under some model of neuronal coupling (Friston, 1994). In particular, gPPI considers a series of equations in which activity in a ROI (pre-defined frontal areas in our case) depends on a specific condition (the ‘psychological’ factor) and on activity in the seed region (striatum here, the ‘physiological’ factor). By solving these equations, it is possible to determine a coefficient that represents task-modulation of effective connectivity^132^. Importantly, task-related changes in effective connectivity are expressed relative to rest, and therefore values closer to 0 reflect a connectivity similar to resting state.

The reward network was defined as following: two regions within the striatum, namely the NAc (BNA regions 223 and 224) and the ventro-medial putamen (BNA regions 225 and 226, left and right respectively), and two frontal areas, namely the anterior cingulate (BNA regions 177, 179, 183 and 178, 180, 184, left and right respectively) and the orbitofrontal cortex within the vmPFC (BNA regions 41, 45, 47, 49, 187 and 42, 46, 48, 50, 188 for left and right respectively). The motor network included the following areas: the dorso-lateral putamen (BNA 229, 230, for left and right respectively), the dorsal caudate (BNA regions 227, 228 for left and right respectively) the medial part of the SMA (BNA regions 9 and 10, left and right respectively) and the part of the M1 associated to upper limb function (BNA regions 57 and 58, left and right respectively). Notably, we considered connectivity in the left and right motor and reward networks regardless of laterality. These ROIs were selected based on the following rationale. First, they are consistent with previous literature on reinforcement learning of motor skills^68,90,133,134^. Second, there is structural and functional evidence for these fronto-striatal connections^135,136^. Third, the frontal areas included in the analyses are well-established hubs of the motor learning (M1 and SMA, see ^12^ for a meta-analysis) and reward networks (vmPFC and ACC, see ^11^ for a meta-analysis). Finally, gPPI was also extracted within a control language network, defined based on the functional atlas described by Shirer et al.(2012)^70^.

## Supplementary material

### 1. Exclusion criteria

- Unable to consent
- Severe neuropsychiatric (e.g., major depression, severe dementia) or unstable systemic diseases (e.g., severe progressive and unstable cancer, life threatening infectious diseases)
- Severe sensory or cognitive impairment or musculoskeletal dysfunctions prohibiting to understand instructions or to perform the experimental tasks
- Color blindness
- Inability to follow or non-compliance with the procedures of the study
- Contraindications for NIBS or MRI:

- Electronic or ferromagnetic medical implants/device, non-MRI compatible metal implant
- History of seizures
- Medication that significantly interacts with NIBS being benzodiazepines, tricyclic antidepressant and antipsychotics
- Regular use of narcotic drugs
- Left-handedness
- Pregnancy
- Request of not being informed in case of incidental findings
- Concomitant participation in another trial involving probing of neuronal plasticity.

### 2. ContES Checklist

**Table S1.**
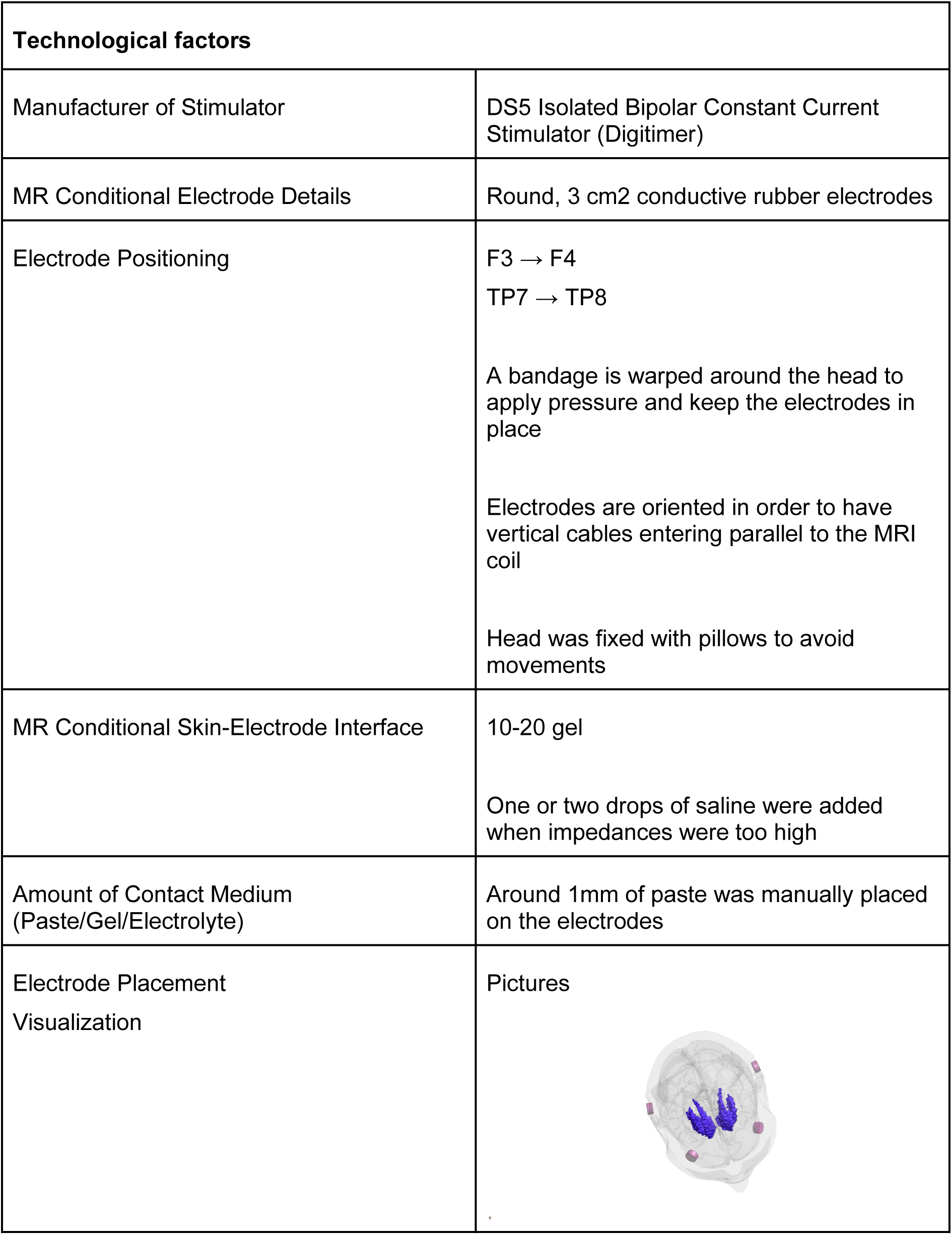
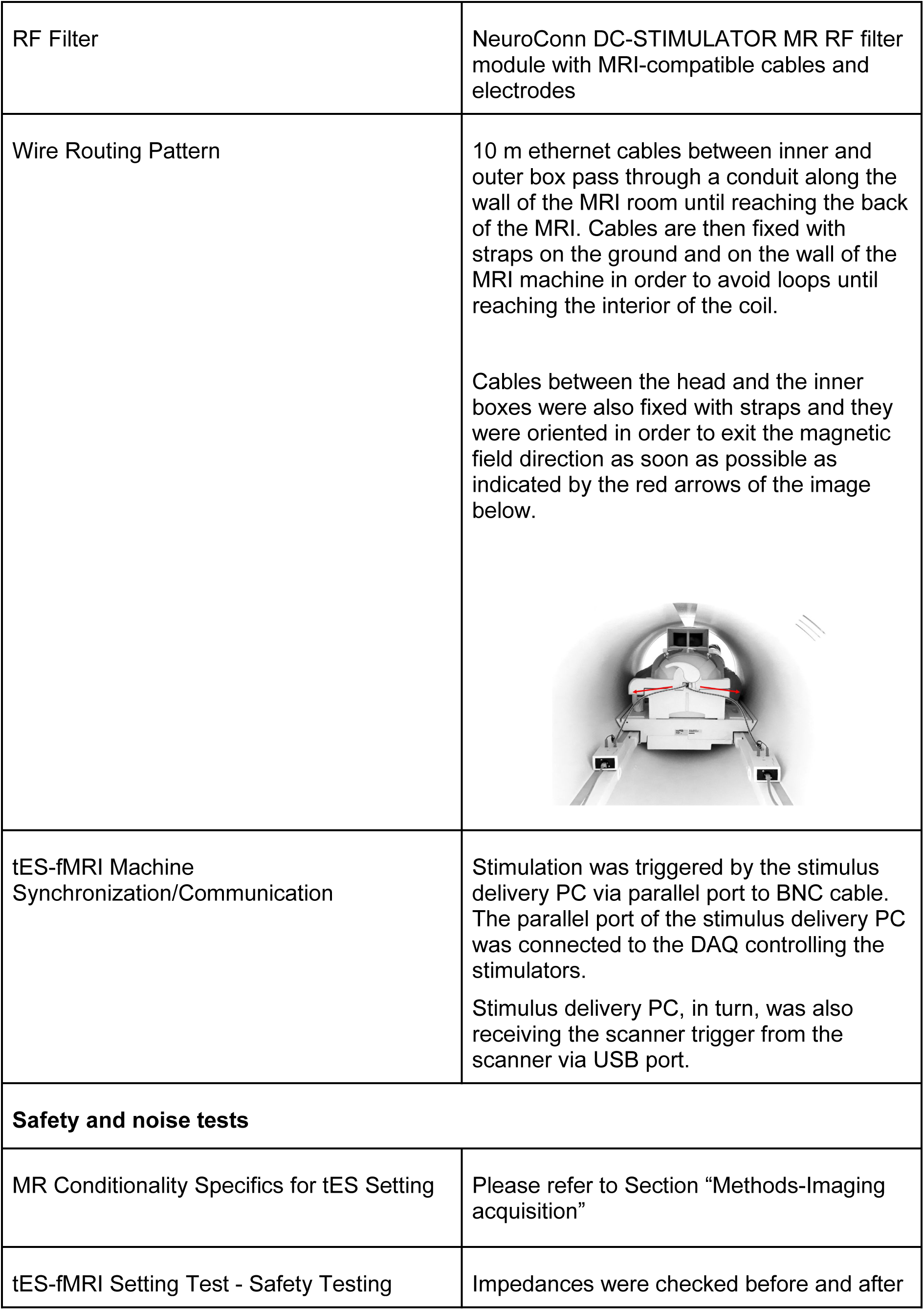
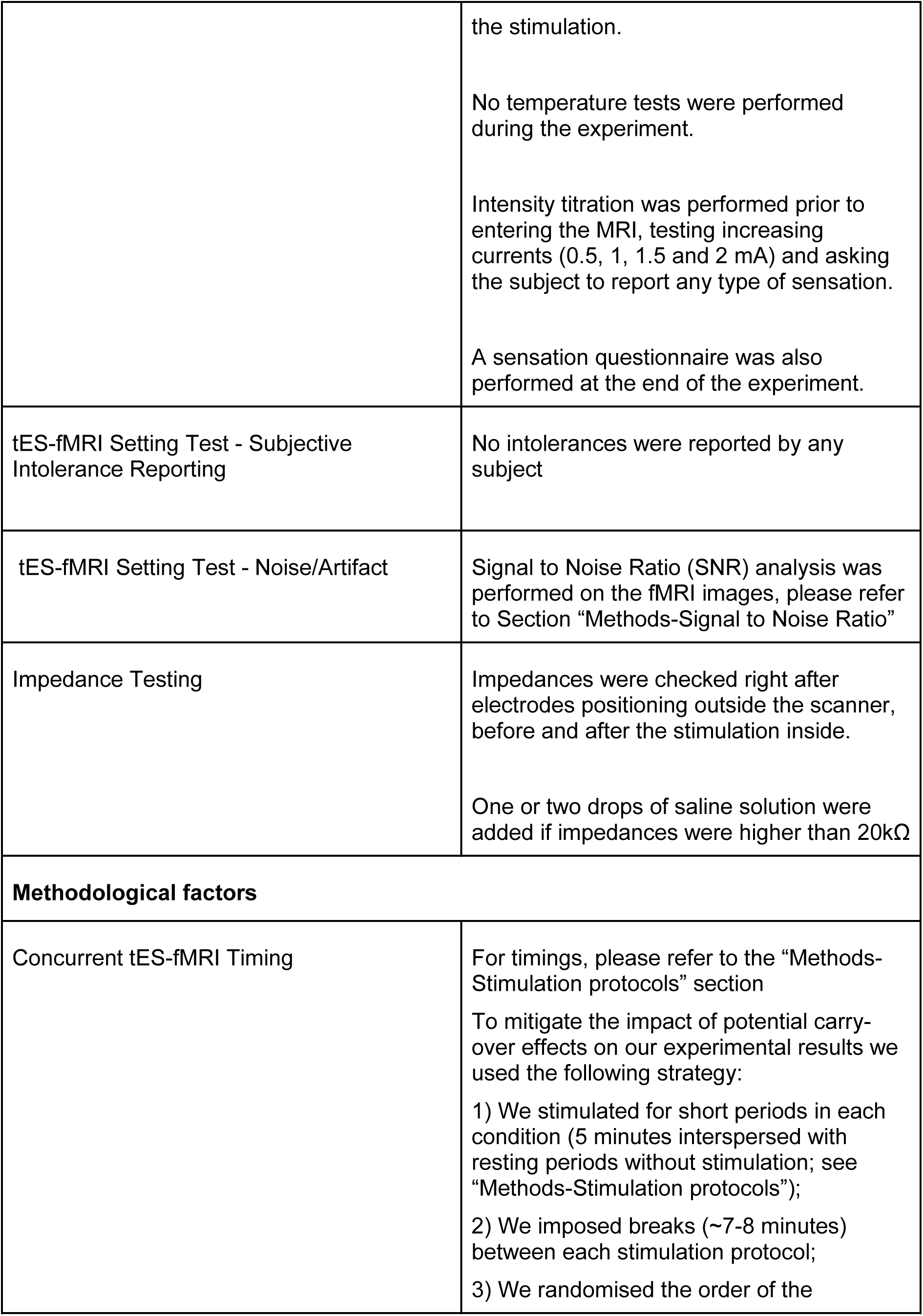
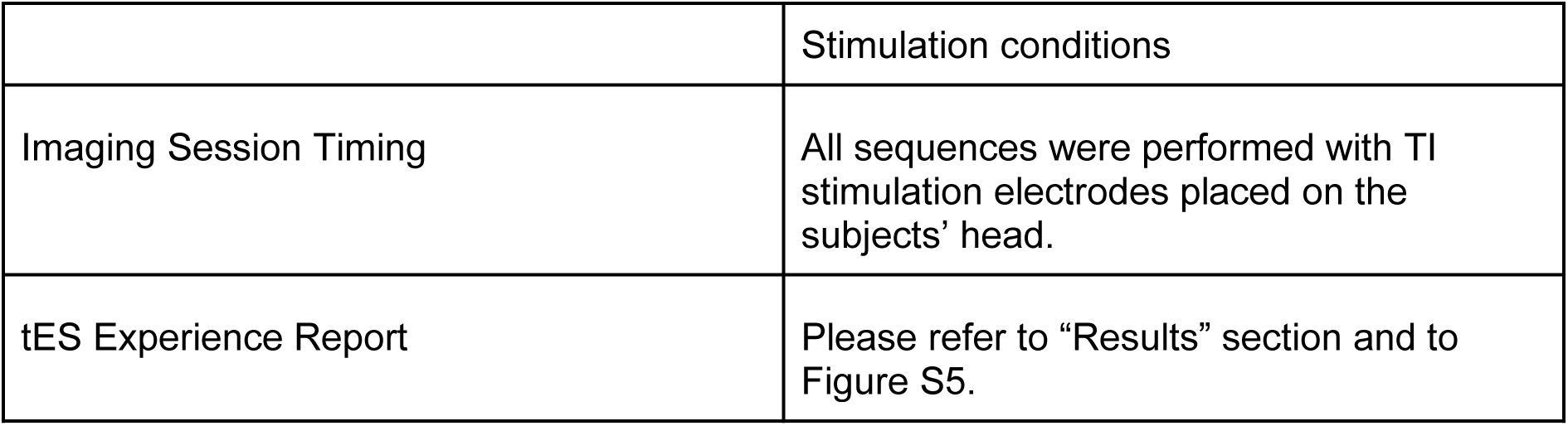
ContES checklist as recommended in Ekhtiari et al., 2022^120^ for concurrent tES-fMRI studies.

### 3. Patterns of motion of the target used in the study

**Figure S1.**
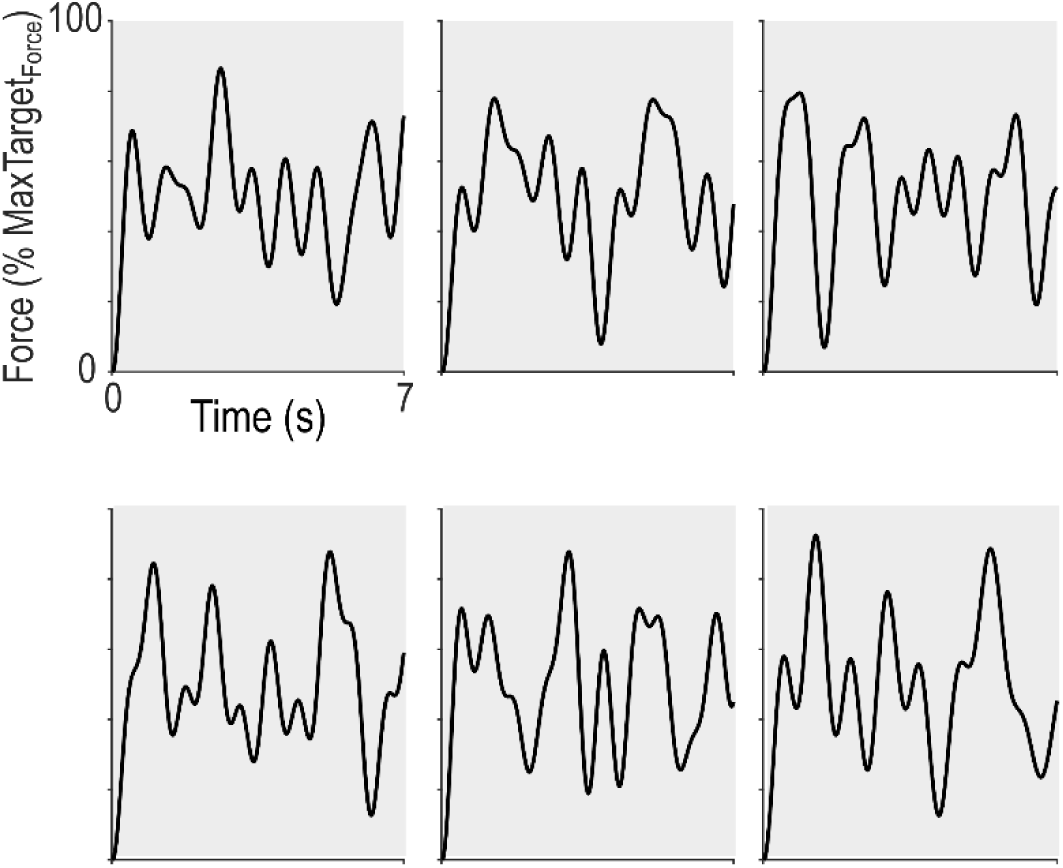
Patterns of motion of the target. For each block of training, participants had to learn a new pattern of motion of the target. The patterns had similar mathematical properties and their relationship to a condition was randomised (see Methods for more details).

### 4. Additional behavioural experiment

To determine the optimal experimental parameters to study reinforcement learning of motor skills, we performed an additional behavioural experiment, in the absence of brain stimulation and imaging. In particular, we tested the relationship between the amount of visual feedback available during Training and the benefits of reinforcement in the force-tracking task. Another group of young healthy participants (n=24; 14 women, 24.2 ± 0.5 years old, independent from the subjects tested in the main experiment) performed blocks of the task with Reinf_ON_ or Reinf_OFF_ and with either full visual feedback or only partial visual feedback (cursor displayed for 35.7% of the total trial duration, as in the main study). Each learning block was composed of 30 trials (vs. 36 trials in the main study) and in addition to real-time closed-loop reinforcement feedback, participants also received endpoint feedback on their overall performance after each trial during Training (i.e., indicating success or failure on the trial). The LMM ran on the Post-training data revealed a significant effect of visual feedback (F_(1,788.33)_=5.90; p=0.015), reinforcement (F_(1,787.87)_=11.64; p<0.001) and a significant interaction between these two factors (F_(1,788.03)_=10.27; p=0.0014, **Figure S2A**). Interestingly, Tukey-corrected post-hoc tests showed that the interaction was due to the fact that while reinforcement did not improve learning when Training was performed with full visual feedback (p=0.88, d=0.014), it induced robust benefits when training with partial visual feedback (p<0.001, d=0.46, **Figure S2B**). This result is in line with previous literature showing that reinforcement feedback is particularly beneficial for motor learning when visual feedback is uncertain^57,62^. Based on the outcome of this additional study, we decided to train participants with partial visual feedback in the present experiment to evaluate the effect of tTIS in a version of the task that yielded significant reinforcement gains. Notably, this work also shows that the effect of reinforcement on motor learning observed in the tTIS_Sham_ and tTIS_20Hz_ conditions (Figure 2) is reproducible.

**Figure S2.**
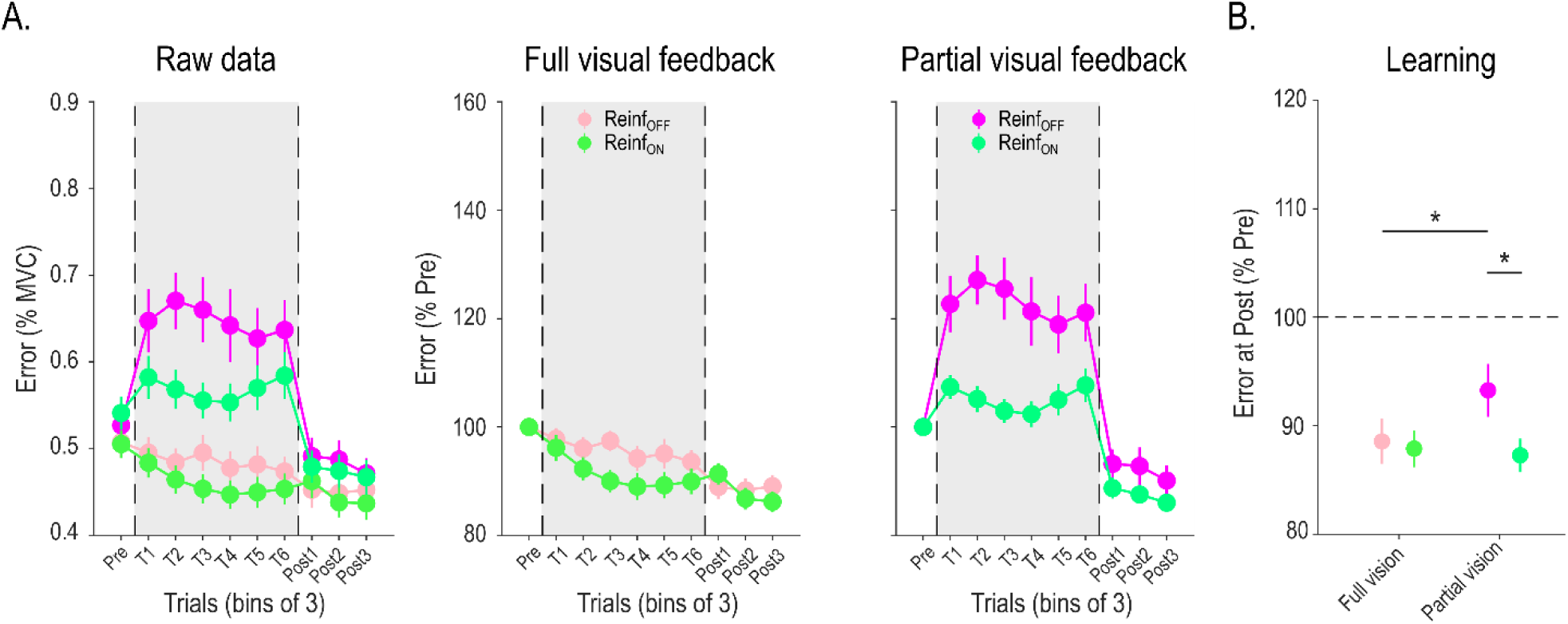
Results of an additional behavioural experiment (n = 24). **A) Motor performance across training.** Raw Error data (expressed in % of Maximum Voluntary Contraction [MVC]) are presented on the left panel for the different experimental conditions in bins of 3 trials. On the right, the two plots represent the Pre-training normalised Error in the Full visual feedback and Partial visual feedback blocks (i.e., cursor displayed 35.7% of the time, as in the main experiment). Note the strong gains in motor performance, especially with partial visual feedback but also the limited improvement of performance during training in this condition. **B) Motor learning.** Averaged Error at Post-training (normalised to Pre-training) in the different experimental conditions are shown, for the subjects included in the analysis (i.e., after outlier detection, n=23). Reduction of Error at Post-training reflects true improvement at tracking the target in Test conditions (in the absence of reinforcement or visual uncertainty). The LMM ran on these data revealed significant a significant effect of reinforcement feedback on learning when training with partial, but not full, visual feedback. *: p<0.05. Data are represented as mean ± SE.

### 5. Evolution of motor performance in the different conditions

The main analysis revealed a general effect of tTIS on motor performance during Training, irrespective of the presence of reinforcement. As a subsequent analysis, we also asked whether the evolution of performance during Training depended on type of striatal stimulation applied. We ran the same LMM as in the main study (see Results) but with the addition of a continuous fixed effect Trial, allowing us to evaluate whether the slope of performance change was different according to tTIS_TYPE_. (**Figure S3**). Indeed, this analysis revealed a significant tTIS_TYPE_ x Trial interaction (F_(2, 3399)_=4.46; p=0.012) that was due to different slopes in the tTIS_Sham_ compared to the tTIS_20Hz_ (p=0.013) and tTIS_80Hz_ (at the trend level, p=0.068) conditions. Evolution of performance in the tTIS_20Hz_ and tTIS_80Hz_ conditions was not different (p=0.81). Notably, this effect could not be explained by differences in initial performance (all p>0.21 when comparing intercepts). Moreover, the effect of tTIS on motor improvement during Training did also not depend on the presence of reinforcement (Reinf_TYPE_, tTIS_TYPE_ and Trial: F_(2, 3399)_=0.51; p=0.60). Overall, this analysis shows that the detrimental effect of striatal tTIS on motor performance is due to an impaired ability to improve performance with practice and further confirms that tTIS did not modulate the ability to use reinforcement feedback during Training.

**Figure S3.**
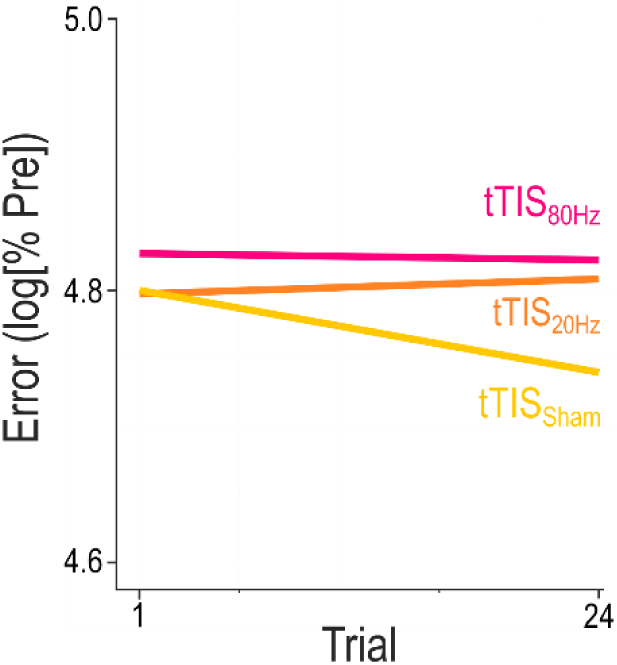
Slopes of performance change during Training in the different stimulation conditions. Modeled performance change for tTIS_Sham_, tTIS_20Hz_ and tTIS_80Hz_ throughout Training. The tTIS_TYPE_ x Trial interaction revealed that performance improved more with tTIS_Sham_ compared to tTIS_20Hz_ and tTIS_80Hz_. Notably this effect was not modulated by the presence of reinforcement and could not be explained by differences in intercepts.

### 6. Effect of visual and reinforcement feedback on motor performance

As a control, we asked whether the tTIS and reinforcement effects reported in Figure 2 depended on the availability of visual information during Training. To do so, we computed the normalised Error for phases with the Cursor_ON_ or Cursor_OFF_ (taking into account a lag of 0.25s, corresponding to the estimated visuo-motor delay in this type of task for young healthy subjects^137^) and analysed these data in a LMM including the factors Reinf_TYPE_, tTIS_TYPE_ and Cursor_TYPE_. As in the main analysis, we confirmed the effect of Reinf_TYPE_ (F_(1, 6872)_=344.87; p<0.001), tTIS_TYPE_ (F_(2,6872)_=28.79; p<0.001) and the absence of interaction between these two factors (F_(2,6875.4)_=0.49; p=0.61, Figure S4). This analysis also revealed a Cursor_TYPE_ effect (F_(2,6875.3)_=49.66; p<0.001) which was due to the fact that the Error was generally higher in the absence visual information on the position of the cursor (d=0.17). Interestingly, there was also a Reinf_TYPE_ x Cursor_TYPE_ interaction (F_(2,6872)_=29.35; p<0.001): while benefits of reinforcement were significant in both the Cursor_ON_ (p<0.001, d=0.32) and Cursor_OFF_ (p<0.001, d=0.58) conditions, the magnitude of the reinforcement-related gains in performance were larger in the Cursor_OFF_ condition (t-test comparing the gains: t_(46)_=2.74, p=0.0086). Moreover, post-hoc tests also revealed that the absence of vision of the cursor was detrimental for performance in the Reinf_OFF_ condition (p<0.001, d=0.30) but not in presence of Reinf_ON_ (p=0.25, d=0.039). Hence, the presence of reinforcement was particularly beneficial when visual information was not available, in line with previous research^57,62^ and also in agreement with the results of our additional experiment (Figure S2). Importantly, the LMM did not reveal any interaction between tTIS_TYPE_ and Cursor_TYPE_ (F_(2,6872)_=0.49; p=0.31) and no triple interaction (F_(2,6872)_=1.53; p=0.22), confirming that striatal tTIS had a global effect on motor performance during Training, which did not depend on the presence of visual and reinforcement feedback.

**Figure S4.**
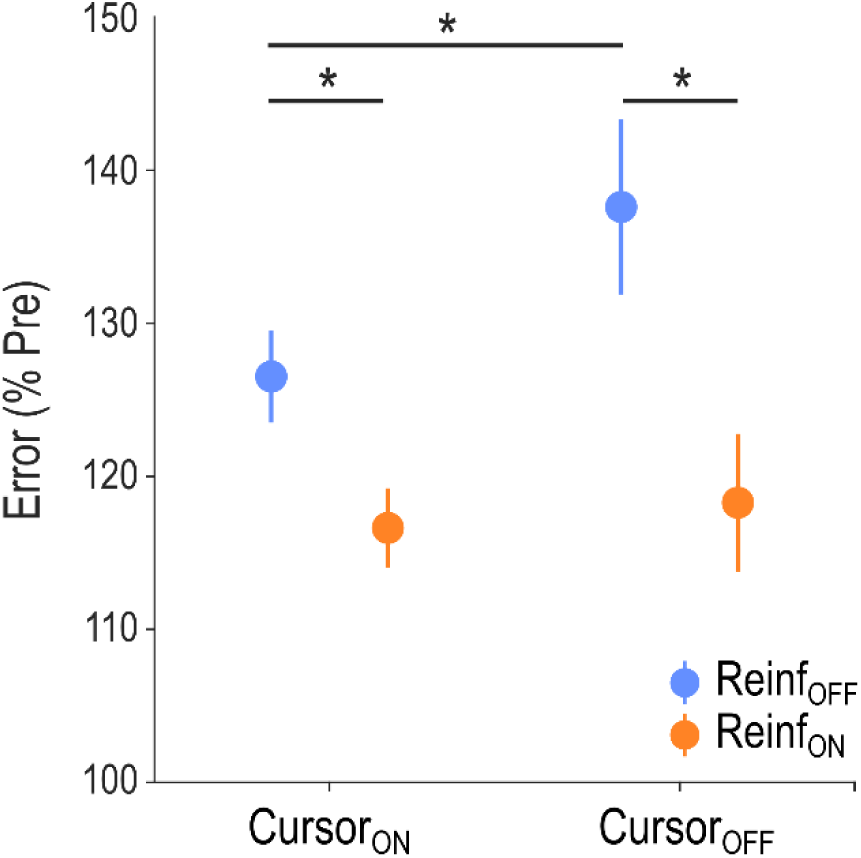
Effect of visual and reinforcement feedback on motor performance. Pre-training normalised Error depending on the presence of the cursor (Cursor_ON_ or Cursor_OFF_), and the presence of reinforcement feedback during Training. The significant Reinf_TYPE_ x Cursor_TYPE_ interaction was related to the fact that the benefits of reinforcement were stronger when visual information was not available. Notably, this analysis takes into account a visuo-motor delay of 0.25s, as previously reported during a similar task (Lam and Zenon, 2021). *: p<0.05. Data are represented as mean ± SE.

### 7. Control analyses of behavioural data

#### Pre-training performance

In order to verify that our main behavioural results were not influenced by potential differences in initial performance between conditions despite randomisation, we analysed the Error at Pre-training between conditions. We did not find any tTIS_TYPE_ (F_(2,519.15)_=1.64; p=0.20) or tTIS_TYPE_ x Reinf_TYPE_ effect (F_(2,519.99)_=1.08; p=0.34), suggesting that the main behavioural results could not be accounted for by differences in initial performance between conditions. However, the LMM did reveal a Reinf_TYPE_ effect (F_(1,519.15)_=12.47; p<0.001), that was due to the fact that Pre-training performance was generally better in Reinf_OFF_ blocks. This effect, which was opposite to our learning results (generally better learning with Reinf_ON_), may be related to an expectancy effect stemming from the repetitive structure of the reinforcement conditions (see Methods). However, the absence of interaction with tTIS_TYPE_ is strongly suggestive that this effect did not drive any of the main findings. Put together, these data provide confidence that the differential effects of striatal tTIS on motor learning depending on the presence of reinforcement were not the result of different initial performance between conditions.

#### Success rate

Overall, the amount of positive reinforcement (i.e., when the target was green) averaged 52.78 +/- 0.42% and was comparable across tTIS_TYPES_ (F_(2,1702)_=0.17; p=0.84), suggesting that the closed-loop reinforcement schedule was successful at providing similar reinforcement feedback despite differences in performance between conditions. Hence, different success rates during training cannot explain the effect of the different striatal tTIS conditions on motor learning.

#### Frequency of flashing

Analysis of the frequency of flashing in the different conditions did not reveal any effect of tTIS_TYPE_ (F_(2,3283)_=0.85; p=0.43) nor any Reinf_TYPE_ x tTIS_TYPE_ interaction (F_(2,3283)_=0.19; p=0.82), suggesting that the behavioural effects of tTIS could not be explained by a visual confound. However, this analysis did reveal a Reinf_TYPE_ effect (F_(1,3283)_=33.62; p<0.001) which was due to the fact that the average frequency in the Reinf_OFF_ condition (4.28 ± 0.097 Hz) was slightly but significantly higher than with Reinf_ON_ (4.08 ± 0.098 Hz; F_(1,3283)_=33.62; p<0.001). Notably, in absolute terms, this difference represented only a difference of 1.4 change of color over the whole 7 s trial, which we think is unlikely to explain the improvement of performance in the Reinf_ON_ condition.

#### Order of the reinforcement conditions

Previous exposure to reinforcement feedback may improve subsequent learning through reinforcement^138^. Thanks to our randomisation procedure, the previous exposure to the Reinf_ON_ condition was equally counterbalanced in all stimulation conditions, and should therefore not influence our main results. Still, we performed an analysis to specifically investigate the effect of the previous exposure to Reinf_ON_. To do so, we split the participants depending on whether they experienced Reinf_ON_ or Reinf_OFF_ first (12 subjects per group) and performed a new LMM on the Post-training data with the addition of a categorical factor Group_TYPE_. In particular, if the previous exposure to the Reinf_ON_ condition influenced following learning with reinforcement, we would expect to see a Group_TYPE_ x Reinf_TYPE_ interaction. The analysis did not indicate any Group_TYPE_ effect on learning (F_(1,21.96)_=0.35; p=0.56), neither did it reveal a Group_TYPE_ x Reinf_TYPE_ (F_(1,4)_=0.72; p=0.44), or a triple interaction with tTIS_TYPE_ (F_(2,1105.06)_=1.75; p=0.17). Overall, this analysis suggests that the order of the exposure to the reinforcement condition did not influence the present findings.

### 8. Blinding integrity and tTIS-evoked sensations

**Figure S5.**
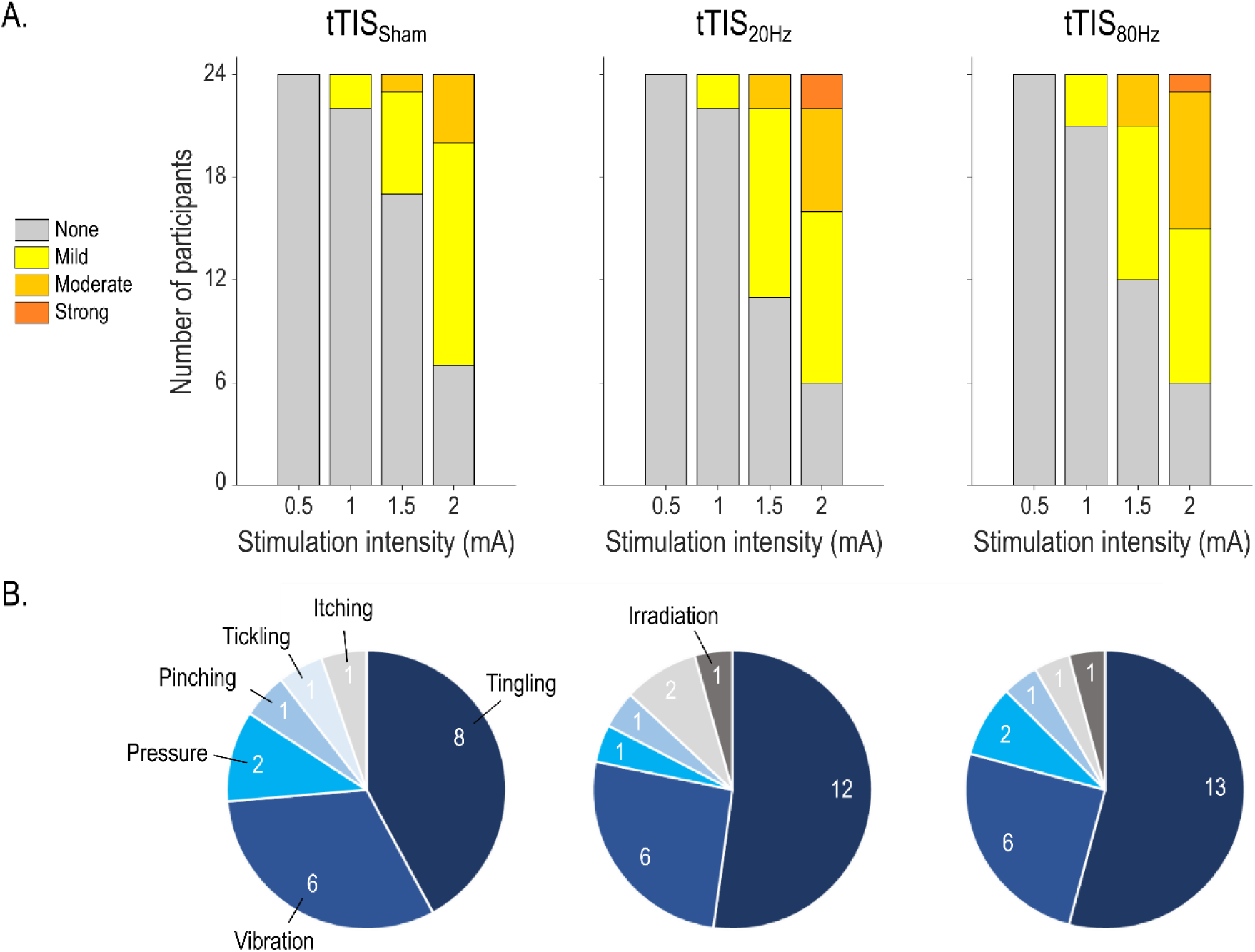
tTIS-related sensations. **A) Magnitude of tTIS-related sensations.** Magnitude of sensations reported before the experiment for current amplitudes ranging from 0.5 to 2 mA for each tTIS_TYPE_. The current amplitude used in the present experiment was 2 mA. **B) Types of tTIS-related sensations.** Type of sensations as described by the participants, at 2 mA. Note that subjects were allowed to describe their sensations with up to two different words.

### 9. Brain activity during reinforcement motor learning

**Figure S6.**
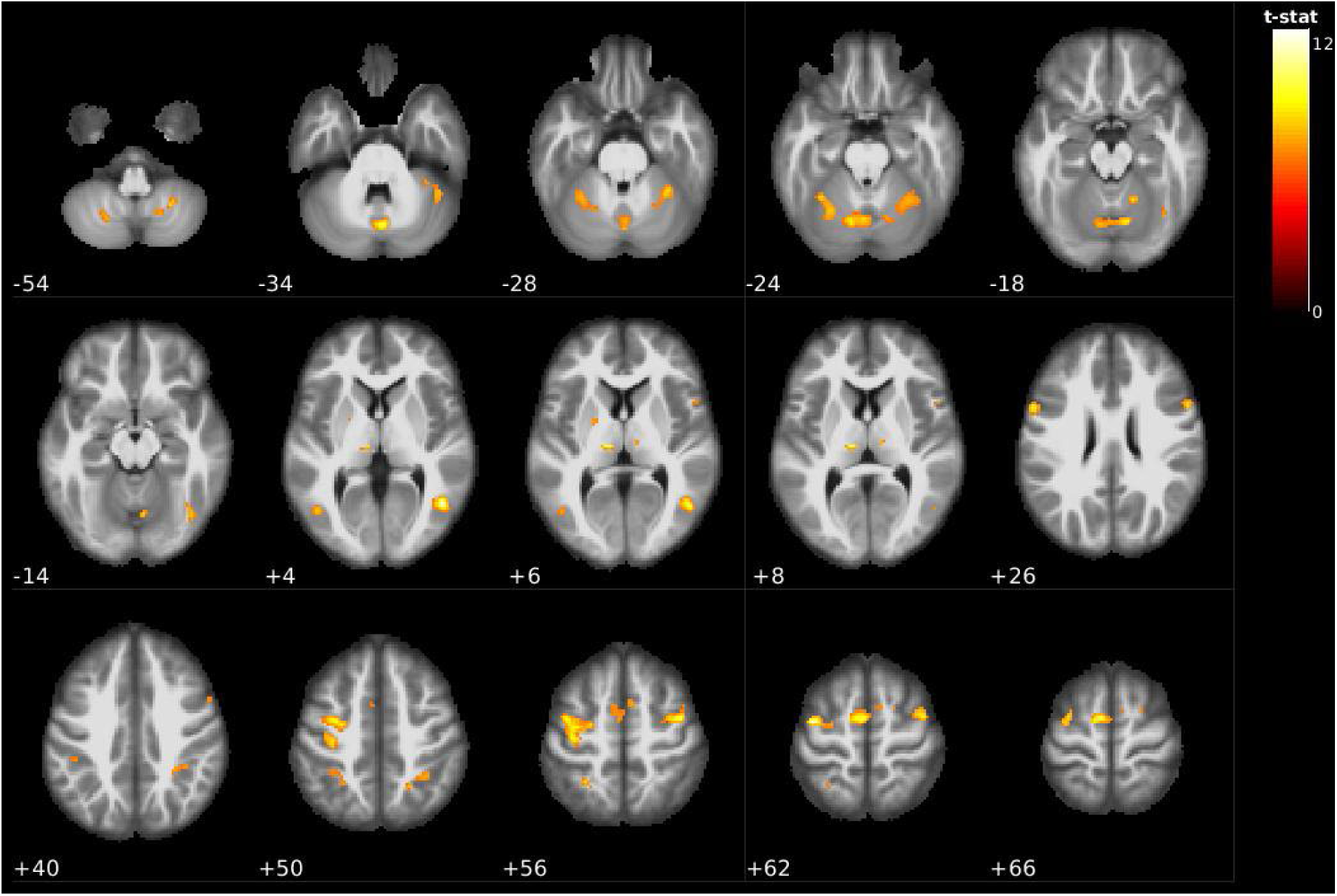
Whole-brain activity during reinforcement motor learning. Activation maps for the contrast task>rest in the tTIS_Sham_, Reinf_ON_ condition showing activation of key areas of the reinforcement motor learning network including the putamen, thalamus, cerebellum and sensorimotor network, especially on the left side. Significant clusters are shown for corrected voxel-wise family wise error (FWE), p=0.05, and corrected cluster-based false discovery rate (FDR), p=0.05.

**Table S2:** Significant clusters and the respective local maxima in the tTIS_Sham_, Reinf_ON_ condition. Related to Figure S6. Regions were identified with the Automated Anatomical Labelling atlas 3 (AAL3^139^). Significant clusters were selected for corrected voxel-wise family wise error (FWE), p=0.05, and corrected cluster-based false discovery rate (FDR), p=0.05.

| Cluster-level |  |  |  | Peak-level |  |  |  |  | x | y | z | Region |
| --- | --- | --- | --- | --- | --- | --- | --- | --- | --- | --- | --- | --- |
| pFWE-corr | qFDR-corr | kE | P<sub>uncorr</sub> | pFWE-corr | qFDR-corr | T | (Z<sub>E</sub>) | P<sub>uncorr</sub> |  |  |  |  |
| <0.001 | <0.001 | 135 | <0.001 | <0.001 | 0.005 | 12.63 | 6.84 | <0.001 | 46 | -62 | 4 | Temporal_Mid_R |
| <0.001 | <0.001 | 523 | <0.001 | <0.001 | 0.005 | 12.32 | 6.77 | <0.001 | -40 | -8 | 62 | Precentral_L |
|  |  |  |  | <0.001 | 0.021 | 10.62 | 6.33 | <0.001 | -34 | -6 | 52 | Postcentral_L |
|  |  |  |  | <0.001 | 0.021 | 10.43 | 6.28 | <0.001 | -36 | -20 | 54 | Precentral_L |
| <0.001 | <0.001 | 335 | <0.001 | <0.001 | 0.018 | 11.08 | 6.46 | <0.001 | -8 | -6 | 64 | Supp_Motor_Area_L |
|  |  |  |  | 0.003 | 0.145 | 8.21 | 5.56 | <0.001 | 6 | 6 | 58 | Supp_Motor_Area_R |
|  |  |  |  | 0.003 | 0.145 | 8.20 | 5.55 | <0.001 | -4 | -2 | 54 | Supp_Motor_Area_L |
| <0.001 | <0.001 | 44 | <0.001 | <0.001 | 0.021 | 10.65 | 6.34 | <0.001 | -10 | -20 | 6 | Thal_IL_L |
| <0.001 | <0.001 | 162 | <0.001 | <0.001 | 0.021 | 10.36 | 6.26 | <0.001 | 42 | -6 | 56 | Frontal_Mid_2_R |
|  |  |  |  | <0.001 | 0.042 | 9.48 | 5.99 | <0.001 | 34 | -4 | 58 | Frontal_Sup_2_R |
| <0.001 | <0.001 | 175 | <0.001 | <0.001 | 0.021 | 10.27 | 6.23 | <0.001 | -58 | 10 | 28 | Precentral_L |
|  |  |  |  | <0.001 | 0.037 | 9.60 | 6.03 | <0.001 | -56 | 8 | 20 | Frontal_Inf_Oper_L |
|  |  |  |  | 0.019 | 0.490 | 7.32 | 5.21 | <0.001 | -48 | 2 | 16 | Rolandic_Oper_L |
| <0.001 | <0.001 | 601 | <0.001 | <0.001 | 0.024 | 10.06 | 6.17 | <0.001 | 2 | -74 | -34 | Vermis_7 |
|  |  |  |  | <0.001 | 0.025 | 9.99 | 6.15 | <0.001 | -12 | -70 | -22 | Cerebellum_6_L |
|  |  |  |  | <0.001 | 0.027 | 9.88 | 6.12 | <0.001 | 12 | -70 | -20 | Cerebellum_6_R |
| <0.001 | <0.001 | 82 | <0.001 | <0.001 | 0.070 | 9.14 | 5.88 | <0.001 | 56 | 10 | 26 | Frontal_Inf_Oper_R |
|  |  |  |  | 0.006 | 0.234 | 7.86 | 5.42 | <0.001 | 56 | 10 | 38 | Precentral_R |
| <0.001 | <0.001 | 141 | <0.001 | 0.001 | 0.092 | 8.89 | 5.80 | <0.001 | -34 | -52 | -24 | Cerebellum_6_L |
|  |  |  |  | 0.002 | 0.117 | 8.47 | 5.65 | <0.001 | -28 | -62 | -24 | Cerebellum_6_L |
| <0.001 | <0.001 | 76 | <0.001 | 0.001 | 0.092 | 8.87 | 5.79 | <0.001 | -28 | -52 | 56 | Parietal_Sup_L |
|  |  |  |  | 0.011 | 0.341 | 7.57 | 5.31 | <0.001 | -30 | -44 | 48 | Parietal_Inf_L |
| <0.001 | <0.001 | 200 | <0.001 | 0.001 | 0.092 | 8.77 | 5.76 | <0.001 | 32 | -48 | -28 | Cerebellum_6_R |
|  |  |  |  | 0.013 | 0.382 | 7.49 | 5.28 | <0.001 | 34 | -40 | -34 | Cerebellum_6_R |
| <0.001 | <0.001 | 36 | <0.001 | 0.001 | 0.092 | 8.73 | 5.74 | <0.001 | 16 | -54 | -18 | Cerebellum_4_5_R |
| <0.001 | <0.001 | 28 | <0.001 | 0.001 | 0.101 | 8.63 | 5.71 | <0.001 | 26 | -58 | -54 | Cerebellum_8_R |
| <0.001 | <0.001 | 62 | <0.001 | 0.001 | 0.113 | 8.51 | 5.67 | <0.001 | 38 | -62 | -16 | Fusiform_R |
|  |  |  |  | 0.002 | 0.117 | 8.45 | 5.64 | <0.001 | 42 | -72 | -12 | Occipital_Inf_R |
| <0.001 | <0.001 | 21 | <0.001 | 0.002 | 0.117 | 8.41 | 5.63 | <0.001 | -46 | -68 | 4 | Occipital_Mid_L |
| <0.001 | <0.001 | 141 | <0.001 | 0.002 | 0.130 | 8.33 | 5.60 | <0.001 | 22 | -56 | 50 | Location not in atlas |
|  |  |  |  | 0.002 | 0.130 | 8.30 | 5.59 | <0.001 | 30 | -48 | 48 | Parietal_Sup_R |
|  |  |  |  | 0.007 | 0.266 | 7.76 | 5.39 | <0.001 | 36 | -40 | 42 | SupraMarginal_R |
| <0.001 | <0.001 | 29 | <0.001 | 0.004 | 0.170 | 8.09 | 5.51 | <0.001 | 44 | -50 | -34 | Cerebellum_Crus_1_R |
| <0.001 | <0.001 | 59 | <0.001 | 0.004 | 0.178 | 8.04 | 5.49 | <0.001 | -22 | -66 | -52 | Cerebellum_8_L |
| <0.001 | 0.006 | 12 | 0.003 | 0.004 | 0.190 | 7.99 | 5.47 | <0.001 | 10 | -16 | 8 | Thal MDI_R |
| 0.001 | 0.043 | 6 | 0.028 | 0.009 | 0.319 | 7.63 | 5.33 | <0.001 | -22 | -2 | 6 | Putamen_L |
| <0.001 | <0.001 | 34 | <0.001 | 0.009 | 0.319 | 7.63 | 5.33 | <0.001 | 18 | -64 | -54 | Cerebellum_8_R |
| 0.001 | 0.300 | 7 | 0.019 | 0.023 | 0.545 | 7.23 | 5.17 | <0.001 | 20 | 2 | 62 | Frontal_Sup_2_R |
| 0.001 | 0.030 | 7 | 0.019 | 0.024 | 0.560 | 7.21 | 5.16 | <0.001 | 52 | 12 | 8 | Frontal_Inf_Oper_R |
| 0.001 | 0.030 | 7 | 0.019 | 0.025 | 0.568 | 7.19 | 5.16 | <0.001 | -44 | -36 | 40 | Parietal_Inf_L |

### 10. Correlation between effect of tTIS_80Hz_ on reinforcement motor learning and modulation of whole-brain activity

**Table S3.**
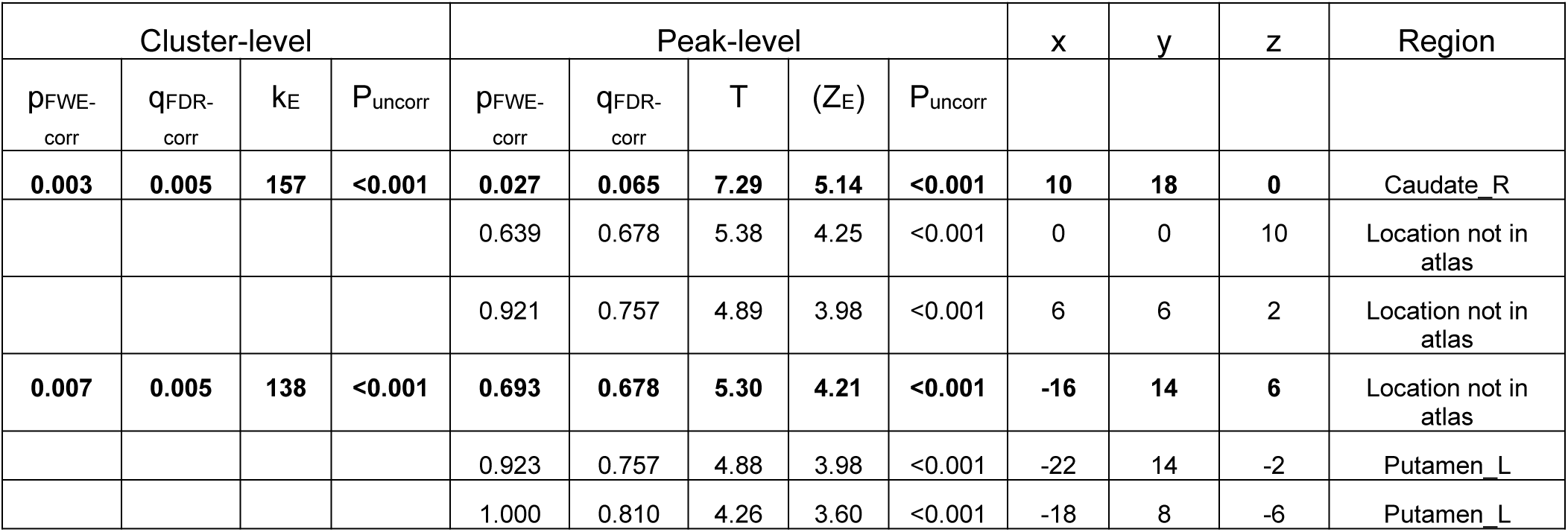
Significant clusters for the correlation between the behavioural and neural effects of tTIS_80Hz_ (vs. tTIS_20Hz_). Related to Figure 3B. Two significant clusters were found with several local maxima. Notably, the left cluster also encompassed a portion of the left caudate (related to Figure 3). Regions were identified with the Automated Anatomical Labelling atlas 3 (AAL3^139^). Significant clusters were selected for uncorrected voxel-wise family wise error (FWE), p=0.001, and corrected cluster-based false discovery rate (FDR), p=0.05.

### 11. Control analysis on striatum to frontal cortex effective connectivity

The connectivity analysis showed that tTIS_80Hz_, but not tTIS_20Hz_, increased striatum to frontal effective connectivity and that this effect depended on the type of network considered (reward vs. motor) and on the presence of reinforcement (Figure 4). In this analysis we considered effective connectivity between the motor striatum and M1 and SMA for the motor network and the limbic striatum with ACC and vmPFC for the reward network, based on a large body of literature^11,12,135,136^ (see Methods for a detailed justification of the ROIs). To verify whether our results depended on the specific frontal ROIs included in the analysis, we performed a new analysis. More specifically, we decomposed connectivity in each network for each frontal cortical area (M1 and SMA in the motor network and ACC and vmPFC in the reward network) and ran two separate LMMs on each network with tTIS_TYPE_, Reinf_TYPE_ as well as ROI_TYPE_ (M1 or SMA for the LMM run on the motor network and ACC or vmPFC for the reward network) as fixed effects. Consistent with our initial findings, we found effects of tTIS_TYPE_ on both LMMs (motor network: F_(2,1089.7)_=3.12; p=0.044 and reward network: F_(2,1112)_=6.78; p=0.0012). Moreover, there was a significant tTIS_TYPE_ x Reinf_TYPE_ interaction in the motor network (F_(2,1112)_=3.36; p=0.035), which was at the trend level in the reward network (F_(2,1113.8)_=2.37; p=0.094). Most importantly, these effects were not modulated by ROI_TYPE_ in any network (tTIS_TYPE_ x Reinf_TYPE_ x ROI_TYPE_ in motor network: F_(2,1112)_=0.83; p=0.44, in reward network: F_(2,1112)_=0.61; p=0.54). This analysis suggests that the main connectivity findings were not influenced by the specific frontal ROIs considered in the analysis.

### 12. Relationship between the neural and behavioural effects of tTIS_80Hz_ and impulsivity

Characterising individual factors that influence responsiveness to brain stimulation is an important line of research both for fundamental neuroscience but also to determine profiles of responders for future clinical translation. Based on previous literature linking striatal gamma oscillatory mechanisms and impulsivity^72^, we explored the possibility that impulsivity influences responsiveness to striatal tTIS_80Hz_ (**Figure S7**).

First, we exploited the BOLD data and asked if inter-individual variability in the neural effects of tTIS_80Hz_ during reinforcement motor learning (i.e., in the Reinf_ON_ condition) was related to impulsivity at the whole-brain level. Impulsivity was evaluated by a well-established independent delay-discounting questionnaire performed at the beginning of the experiment^75,76^. Strikingly, this analysis revealed that impulsivity was associated to the effect of tTIS_80Hz_ (with respect to tTIS_20Hz_) specifically in the left caudate nucleus (Figure S7A, Table S4). No other clusters were found. As such, the most impulsive participants exhibited an increase of left caudate activity with tTIS_80Hz_ (compared to tTIS_20Hz_) while the least impulsive ones rather presented a decrease of BOLD signal, consistent with the idea that impulsivity modulates the neuronal responsiveness to tTIS (R^2^=0.47; p<0.001; Figure S7B). No significant clusters of correlation were found for the tTIS_80Hz_ – tTIS_Sham_ contrast, neither for the control tTIS_20Hz_ -tTIS_Sham_ contrast. Hence, this analysis suggests that the effect of tTIS_80Hz_ on caudate activity depends on participants’ impulsivity.

As a second step, we aimed at evaluating the association between impulsivity and the increased striatum to motor cortex connectivity observed with tTIS_80Hz_, in the presence of reinforcement. Notably, such pattern of increased connectivity in fronto-striatal circuits has been described as a pathophysiological mechanism in multiple neuro-psychiatric disorders involving impulsivity^101–104^. Hence, we first asked if striatum to motor cortex connectivity was related to impulsivity during reinforcement motor learning in the absence of stimulation (i.e., in the tTIS_Sham_ condition). Indeed, we found a significant positive relationship between impulsivity and striatum to motor cortex connectivity (robust linear regression: R^2^=0.10; p=0.0038), in line with previous results^101–104^. Then, we evaluated whether the increase of connectivity observed with tTIS_80Hz_ in the Reinf_ON_ condition (Figure 4A) could be related to impulsivity. Indeed, we found that the effect of tTIS_80Hz_ on connectivity was negatively correlated to impulsivity both when contrasting tTIS_80Hz_ with tTIS_Sham_ (R^2^=0.19; p=0.043, Figure S7C, left) and with tTIS_20Hz_ (R^2^=0.28; p=0.021, Figure S7C, middle): participants with the largest increase in connectivity with tTIS_80Hz_ in the Reinf_ON_ condition were also the least impulsive ones. Such correlation was absent when contrasting tTIS_20Hz_ and tTIS_Sham_ (R^2^=0.0031; p=0.31, Figure S7C, right), but also when considering the same contrasts in the reward instead of the motor network (p=0.93 and p=0.86 for the tTIS_80Hz_-tTIS_Sham_ and tTIS_80Hz_-tTIS_20Hz_ contrasts, respectively). Hence, striatum to motor cortex effective connectivity during the task was positively correlated to impulsivity, but the change in connectivity induced by tTIS_80Hz_ was rather negatively associated with impulsivity. This may be due to a ceiling effect in the most impulsive participants: exhibiting initially high levels of connectivity may leave less room for further modulation by tTIS_80Hz_. These results suggest that inter-individual variability in impulsivity might influence neural responses to striatal tTIS_80Hz_.

**Figure S7.**
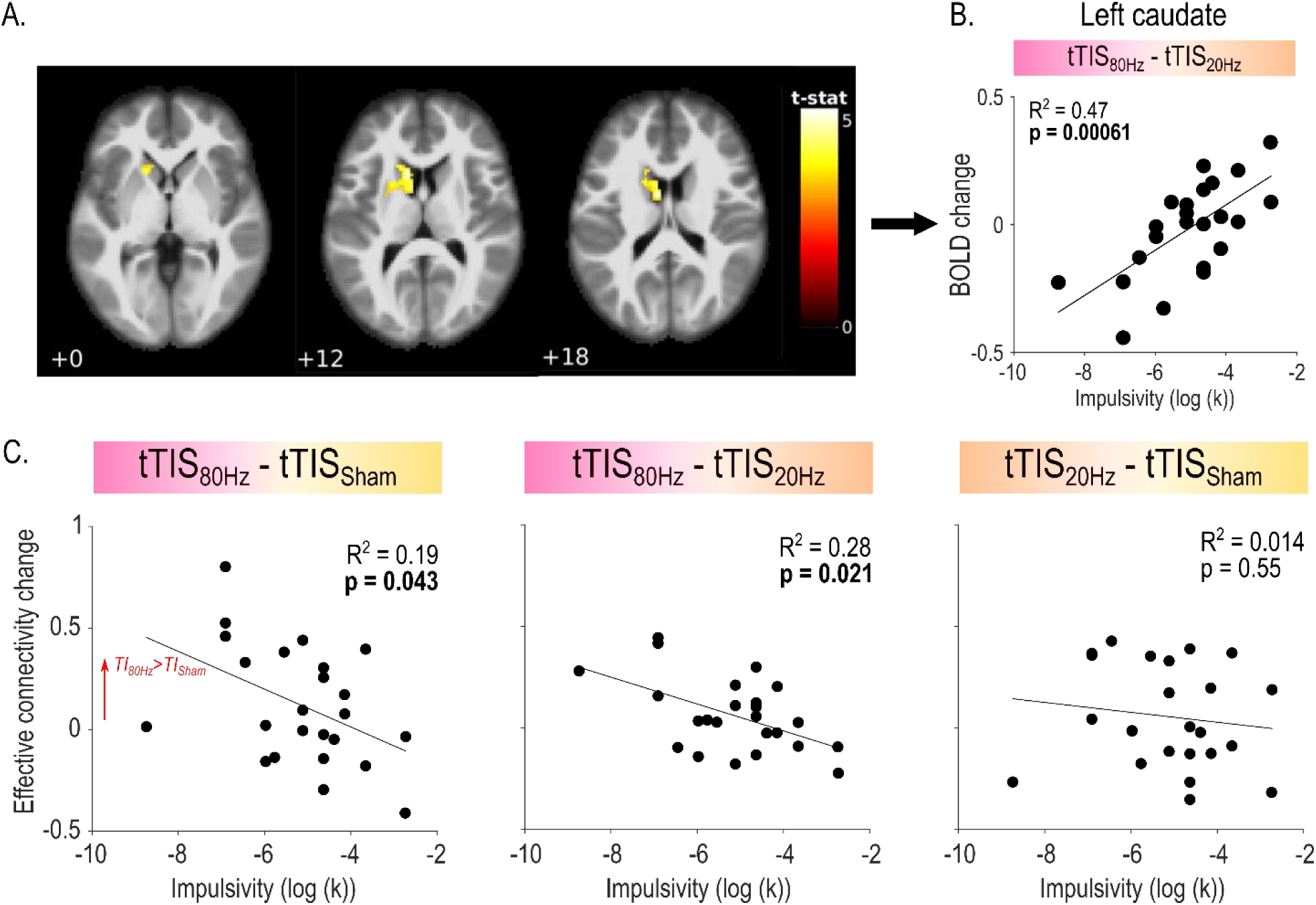
Relationship between impulsivity and the neural effects of tTIS_80Hz_. **A) Whole-brain correlation between the neural effects of tTIS_80Hz_ (with respect to tTIS_20Hz_) and impulsivity.** Correlation between tTIS-related modulation of striatal activity (tTIS_80Hz_ – tTIS_20Hz_) during reinforcement motor learning (Reinf_ON_) and individual impulsivity levels. A single significant cluster of correlation was found in left caudate (uncorrected voxel-wise FWE: p=0.001, and corrected cluster-based FDR: p=0.05). **B) Correlation between left caudate activity and impulsivity.** A positive correlation was found showing that participants with higher levels of impulsivity exhibited stronger activation of the left caudate in the tTIS_80Hz_ (with respect to tTIS_20Hz_). **C) Correlations between impulsivity and tTIS-related modulation of effective connectivity.** Impulsivity was associated to the neural effects of tTIS_80Hz_ both when contrasting to tTIS_Sham_ (left) and tTIS_20Hz_ (middle), but was not correlated to the effect of tTIS_20Hz_ (right).

**Table S4.** Significant clusters for the correlation between impulsivity and effects of tTIS_80Hz_ on BOLD activity (vs. tTIS_20Hz_). Related to Figure S7A. One significant cluster encompassing the left caudate nucleus was found. Regions were identified with AAL3^139^.

| Cluster-level |  |  |  | Peak-level |  |  |  |  | x | y | z | Region |
| --- | --- | --- | --- | --- | --- | --- | --- | --- | --- | --- | --- | --- |
| pFWE-corr | qFDR-corr | kE | P<sub>uncorr</sub> | pFWE-corr | qFDR-corr | T | (ZE) | P<sub>uncorr</sub> |  |  |  |  |
| <0.001 | <0.001 | 254 | <0.001 | 0.707 | 0.524 | 5.29 | 4.20 | <0.001 | -8 | 0 | 18 | Location not in atlas |
|  |  |  |  | 0.719 | 0.524 | 5.27 | 4.19 | <0.001 | -14 | 16 | 16 | Caudate_L |
|  |  |  |  | 0.971 | 0.620 | 4.72 | 3.88 | <0.001 | -16 | 16 | 0 | Location not in atlas |

As a last step, we verified if impulsivity was also predictive of the behavioural effects of tTIS_80Hz_ on reinforcement motor learning. We did not find any significant correlation between impulsivity and the effect of tTIS_80Hz_ on motor learning (tTIS_80Hz_ – tTIS_Sham_: R^2^=0.098; p=0.17; tTIS_80Hz_ – tTIS_20Hz_: R^2^=0.11; p=0.21). Hence, impulsivity was associated to the neural, but not the behavioural effects of tTIS_80Hz_.

Overall, we found that impulsivity was associated to tTIS_80Hz_-related BOLD changes specifically in the left caudate and to changes of effective connectivity between the motor striatum and motor cortex during reinforcement motor learning. Hence, a possibility is that the differences in endogenous striatal gamma-related activity that have been associated to impulsive behaviour in animal models^72–74^, influence the neural effects of tTIS_80Hz_. If this is the case, impulsivity could constitute a behavioural factor allowing to determine responsiveness to striatal tTIS_80Hz_. Conversely, an interesting avenue for future research could aim at determining whether impulsivity can be modulated by striatal tTIS_80Hz_.

### 13. Imaging quality control

A threshold of 0.5 was chosen to discard subjects showing more than 40% of voxels with framewise displacement FD higher than this threshold. In the current study cohort, no subject exceeded the limit value, thus the whole dataset could be used. Furthermore, successful cleaning of the data was ensured by visual checking the preprocessing results. In particular, good registration between anatomical and functional images and normalization to standard space were checked. Signal to noise ratio analysis showed significantly higher tSNR values underneath the stimulating electrodes (F_(1,1122)_=249.25, p<0.001; **Figure S5**). Moreover, an additional analysis showed that this effect was not influenced by the tTIS_TYPE_ (Sphere_LOCATION_ x tTIS_TYPE_: F(2,1118)=0.0169, p=0.98). This result suggests that the stimulation did not introduce additional noise to the MR images. In summary, all controls confirmed the good quality of the imaging data.

**Figure S8.**
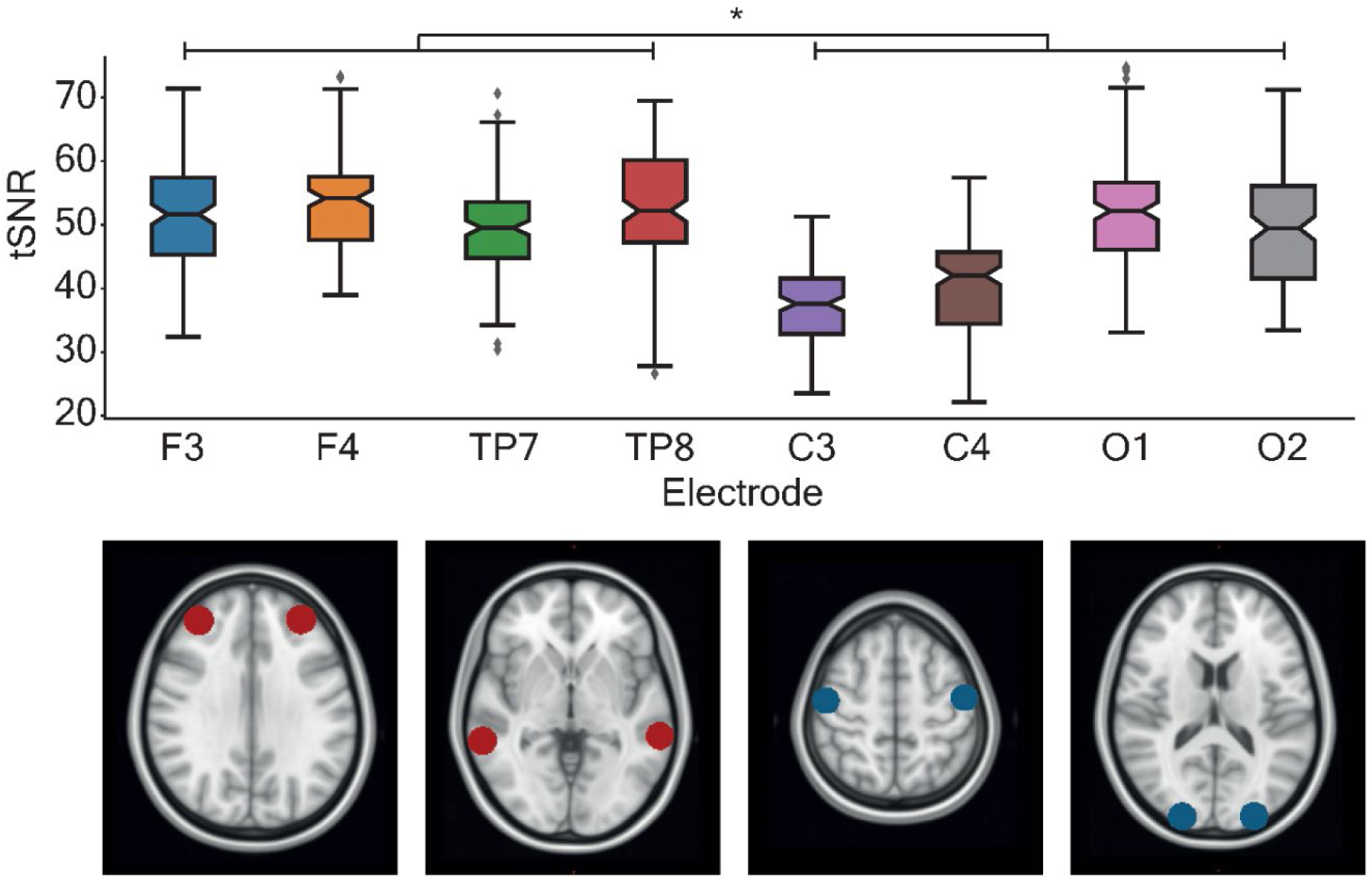
Total signal to noise ratio (tSNR). Total signal to noise ratio investigation. On the top panel, the average tSNR is shown within spheres of 10mm radius underneath the 4 stimulation electrodes (F3, F4, TP7 and TP8) and underneath other 4 locations more distal from the electrodes (C3, C4, O1 and O2). A significant higher tSNR was found underneath the electrodes with respect to the distal locations (F(1,1122)=249.25, p<0.001). This indicates that there was no reduction of the tSNR due to the presence of electrical current. On the bottom panel, the location of the spheres from where the average tSNRs were extracted: F3 and F4 in red in the first image from the left, TP7 and TP8 in red on the second image from the left, C3 and C4 in blue on the third image from the left, O1 and O2 in blue on the forth image from the left.

## Acknowledgements

P.V. was a PhD student supported by the Fund for Research training in Industry and Agriculture (FRIA/FNRS; FC29690), and grants by the Platform for Education and Talent (Gustave Boël - Sofina Fellowships) and Wallonie-Bruxelles International. J.D. was supported by grants from the Belgian FNRS and the Fondation Médicale Reine Elisabeth (FMRE). G.D. was supported by the Belgian FNRS. The research was partially funded by the Novartis Research Foundation—FreeNovation (Basel, CH) to MJW and EN, the Bertarelli Foundation (Catalyst ‘Deep-MCI-T’, Gstaad, CH) to FCH, the SNSF Lead Agency (NiBS-iCog 320030L_197899) to FCH and the Defitech Foundation (Morges, CH) to FCH. We acknowledge access to and expertise of the Neuromodulation and the Neuroimaging facilities of the Human Neuroscience Platform of the Campus Biotech Geneva and of the MRI Platform of the HVS (Sion). We would also like to thank Prof. Leonardo Cohen for insightful feedback on a previous version of this manuscript.

## Data and code availability statement

The datasets generated during the current study and the code used to analyse them are available from the corresponding author on reasonable request.

